# Optogenetic control of beta-carotene bioproduction in yeast across multiple lab-scales

**DOI:** 10.1101/2022.10.31.514479

**Authors:** Sylvain Pouzet, Jessica Cruz-Ramon, Matthias Le Bec, Céline Cordier, Alvaro Banderas, Simon Barral, Benoit Sorre, Sara Castano-Cerezo, Thomas Lautier, Gilles Truan, Pascal Hersen

**Affiliations:** Institut Curie, Université PSL, Sorbonne Université, CNRS UMR168, Laboratoire Physico Chimie Curie, 75005 Paris, France; Toulouse Biotechnology Institute, Université de Toulouse, CNRS, INRAE, INSA, Toulouse, France

**Keywords:** Optogenetics, bioproduction, *Saccharomyces cerevisiae*, beta-carotene, synthetic biology, metabolic engineering

## Abstract

Optogenetics arises as a valuable tool to precisely control genetic circuits in microbial cell factories. Light control holds the promise of optimizing bioproduction methods and maximize yields, but its implementation at different steps of the strain development process and at different culture scales remains challenging. In this study, we aim to control beta-carotene bioproduction using optogenetics in *Saccharomyces cerevisiae* and investigate how its performance translates across culture scales. We built four lab-scale illumination devices, each handling different culture volumes, and each having specific illumination characteristics and cultivating conditions. We evaluated optogenetic activation and beta-carotene production across devices and optimized them both independently. Then, we combined optogenetic induction and beta-carotene production to make a light-inducible beta-carotene producer strain. This was achieved by placing the transcription of the bifunctional lycopene cyclase / phytoene synthase CrtYB under the control of the pC120 optogenetic promoter regulated by the EL222-VP16 light-activated transcription factor, while other carotenogenic enzymes (CrtI, CrtE, tHMG) were expressed constitutively. We show that illumination, culture volume and shaking impact differently optogenetic activation and beta-carotene production across devices. This enabled us to determine the best culture conditions to maximize light-induced beta-carotene production in each of the devices, reaching a content of up to 880 μg/gCDW. Our study exemplifies the stakes of scaling up optogenetics in devices of different lab scales and sheds light on the interplays and potential conflicts between optogenetic control and metabolic pathway efficiency. As a general principle, we propose that it is important to first optimize both components of the system independently, before combining them into optogenetic producing strains to avoid extensive troubleshooting. We anticipate that our results can help designing both strains and devices that could eventually lead to larger scale systems in an effort to bring optogenetics to the industrial scale.

## INTRODUCTION

Advances in bioproduction using yeast and bacteria as microbial cell factories have enabled significant feats in metabolic engineering to be showcased ^1,2^ and allow for the unprecedented production of complex chemicals through more sustainable processes. However, despite impressive progresses, the compound of interest is often produced at relatively low levels^3^. To increase production yields, forward metabolic engineering (rational design and Design-Build-Test-Learn Cycles)^3^ and reverse metabolic engineering (mutagenesis and directed evolution)^4^ have emerged as preferred strategies to increase the carbon flux and redirect more cellular resources towards the production of the compound of interest. These strain engineering strategies are increasingly benefiting from progress in computational approaches (multiomics^5,6^, genome-wide metabolic models^5^, machine learning^7^).

To improve yields, complementary strategies mostly focus at the bioprocess level. There, the key is to best cope with metabolic burden, *i*.*e*., the potential metabolic load and stress created during production, such that growth is least impaired, and productivity maintained. To this end, tools and methods to decouple growth from production have been continuously perfected since the first production of antibiotics from fungi^8^. Inducible systems (usually relying on transcriptional activation) are often employed to control the shift from the growth phase (building up biomass without production nor burden) to the production phase (which focuses on production with minimal growth).

Moving beyond this two-step cultivation strategy, several recent studies showed that controlling growth *versus* production in a dynamic manner can further increase production yields^9,10^. Finely controlling the induction of the metabolic pathways leading to production is critical for such “dynamic regulation”^11^ strategies. The inducer must be reversible, responsive, easy to handle, and cheap. Today, mostly chemical inducers are used. But once injected in large volumes, chemical inducers take time to diffuse uniformly, which leads to potential induction heterogeneity, and cumbersome medium changes are usually required to reverse their action. Besides, metabolizable carbon-sources used as inducers (galactose, methanol, arabinose) exhibit slow response dynamics and are expensive at larger scales. In contrast, the use of light as an inducer (optogenetics) is a very attractive strategy since the responsiveness and its reversibility are instantaneous.

Optogenetics has already been applied to bioproduction, though only in a handful of lab-scale studies. Light has mostly been used to control transcription irreversibly^12^ or reversibly^9,13–15^, to direct the assembly of enzymatic clusters^16^, or to tune the composition of microbial consortia^17^. Development of an optogenetic producer strain at the lab-scale requires the use of specific illumination devices. In this paper, we present and compare four devices that can be used at every step of the strain development process: from 24-well plate systems (OptoBox^18^), to simple small-scale starter cultures (OptoTubes), larger-scale batch cultures (OptoFlasks), and devices that allow multiple-parameter control of the culture conditions (eVOLVER^19^).

We used the EL222 optogenetic system, a single-component light-oxygen-voltage-sensing (LOV)-based light-activated synthetic transcription factor derived from the bacteria *Erythrobacter litoralis*. EL222 offers rapid activation (τ_on_ = 5sec) and deactivation kinetics (τ_off_ = 30sec)^20^ and is an established system in yeast^9^.

Beta-carotene is a terpene known for its characteristic yellow/orange color. Its photochemical and antioxidant properties make this carotenoid a valuable molecule in a wide range of industries, including cosmetics (sun protection), health (antioxidant, dietary supplement as provitamin A), feed and food (health and coloring)^21^. Major sources of beta-carotene are chemical synthesis (enol-ether condensation or Wittig condensation of beta-ionone^22^) and extraction from plants (carrots, oil of palm fruit, and sweet potato) or naturally producing microorganisms (mostly the algae *Dunaliella spp*. and fungus *Blakeslea trispora*)^21^. Beyond its industrial relevance, beta-carotene and other compounds of economic importance (artemisinine^23^, taxol^24^, celastrol^25^) originate from the isoprenoid pathway. Therefore, new improvements obtained using beta-carotene could also benefit the production of other compounds. In addition, production of a colorful compound that can be detected with the naked eye is a particularly convenient output for bioproduction studies. Hence, beta-carotene is often used as a proxy for various proofs-of-concept and genetic systems^26–29^.

In this study, we use the EL222 optogenetic system and the beta-carotene synthesis pathway to explore the importance of different culture parameters at various scales using four illumination devices. The constraints encountered at these different scales of lab culture foreshadow larger-scale potential challenges. Indeed, differences in illumination (light input, medium penetration, and device geometry) and the culture conditions (stirring and shaking, culture volume, gas exchange) between these scales can independently impact both genetic components: the activation of the optogenetic system and the bioproduction by the metabolic pathway. First, we detail the constructions of these four illumination devices and investigate how optogenetics translates across lab scales. Second, we optimize culture conditions to maximize beta-carotene production across devices and identify production constraints. Only then, as a third step, we combine optogenetic control with beta-carotene production. We investigate to what extent this optogenetic control of bioproduction recapitulates constraints identified independently for both genetic components and discuss the relationships between optogenetic activation and the resulting production. In a nutshell, we demonstrate how this three-step approach can help identify optogenetic, metabolic and scaling constraints to define the optimal parameters to drive beta-carotene production in response to light in the budding yeast *Saccharomyces cerevisiae* in multiple lab culture scales.

## RESULTS AND DISCUSSION

### Setting-up optogenetics in different lab-scale culture devices

#### OptoBox

The OptoBox, (*a*.*k*.*a*. The Light-Plate Apparatus^18^ – Fig.1A) was developed to illuminate 24-well imaging plates in a shaking incubator. Each well of the OptoBox contains two interchangeable LEDs connected to a Printed Circuit Board (PCB). Although the two LEDs can be different in order to be compatible with various optogenetic systems, we connected two blue LEDs (461 nm) to maximize light exposure for the EL222 optogenetic system. Programing of the device is particularly easy: the program can be set up online using the user-friendly software Iris, uploaded to an SD card, and plugged into the PCB. The programming consists of assigning each of the two LEDs an arbitrary value between 0 and 4000 (for a given illumination pattern), corresponding, in our case, to an intensity of 0 to 4mW/cm^2^ per LED (Fig.S1). The OptoBox provides a convenient tool to screen various strains or illumination conditions. The 24-well plates generally hold a culture volume of 1 mL per well and can be sealed with aluminum foil to prevent light leaking, culture spilling and evaporation. At the bottom of the well-plate we use (see Methods), lies a 25 μm film that allows for gas exchanges and light penetration.

**Figure 1.**
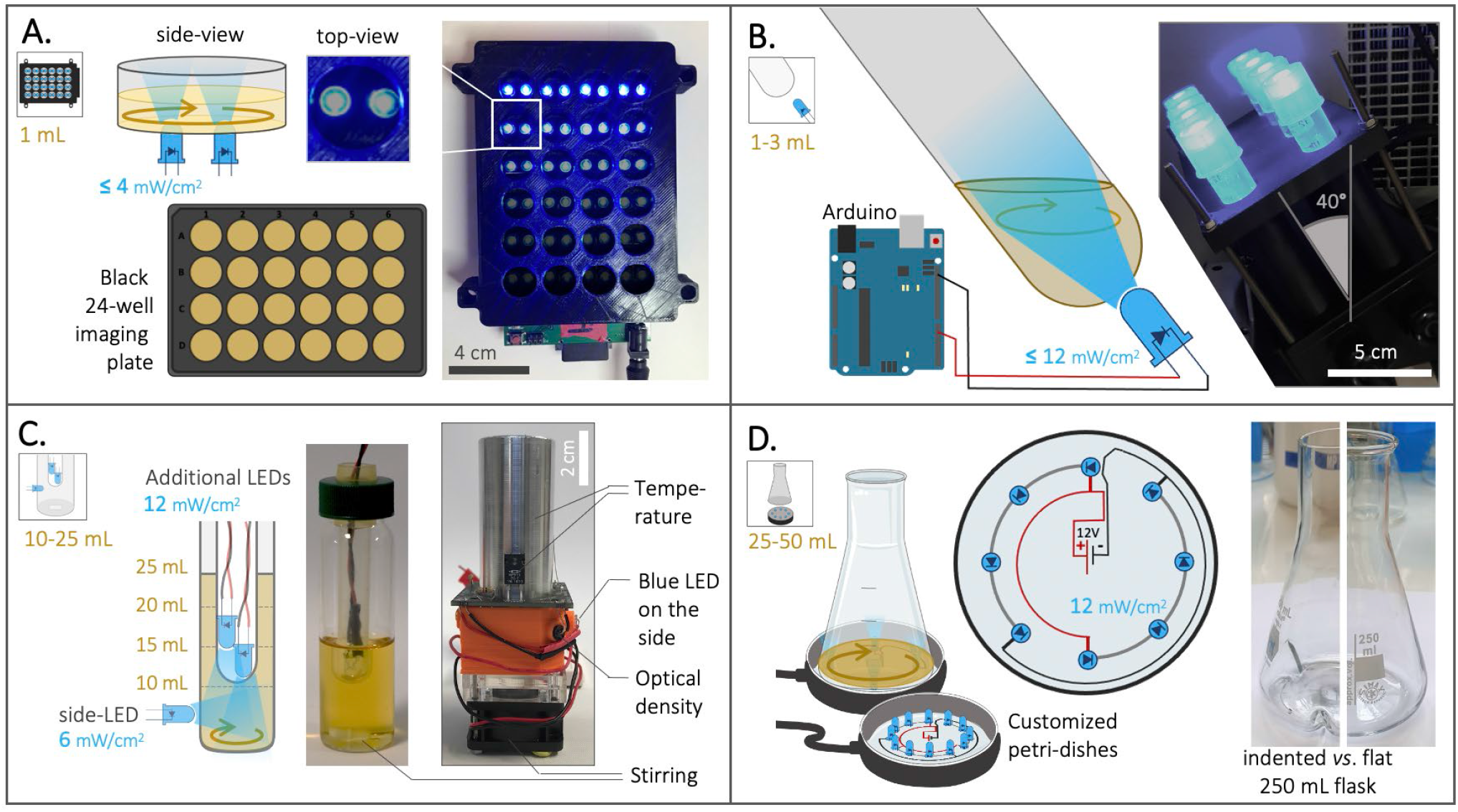
Description of the four optogenetic devices used in this study. **(A)** The OptoBox (adapted from Gerhardt *et al*. 2016^18^) can independently illuminate 1 mL cultures in a 24-well plate placed in a shaking incubator. Two LEDs (0 to 4 mW/cm^2^) illuminate each well from below and can be programmed. **(B)** The OptoTubes are used to illuminate 14 mL tubes (generally 3 mL cultures) using a LED (0 to 12 mW/cm^2^) placed at the bottom of each tube. The OptoTubes can be programmed with an Arduino (0 to 255 u.a.) and be placed in a shaking incubator thanks to a dedicated 3D printed opaque holder. **(C)** The eVOLVER culture platform adapted from Wong *et al*. 2018^19^ uses a DIY “sleeve” (right) where a glass vial (center) can be inserted, all connected to an Arduino. We built 16 of these units, and the temperature, stirring and illumination (via an additional side-LED: 6 mW/cm^2^) can be controlled for each unit, while the growth rate and production of beta-carotene can be monitored. The lid of the glass vial was adapted to accommodate the input of more light using additional LEDs (12 mW/cm^2^ each). **(D)** The OptoFlasks, in which custom-made illumination stands were built to hold different numbers of LEDs (12 mW/cm^2^ each), on top of which the flasks (indented/baffled or flat) are positioned and can hold 25 to 50 mL cultures.

#### OptoTubes

To illuminate simple culture tubes with light, we designed a set of OptoTubes by positioning LEDs at the bottom of 14 mL glass test tubes (Fig. 1B). The LEDs are directly connected to an Arduino’s Pulse-Width Modulation (PWM) pin for intermediate light intensities (from 0 to 255, corresponding to 0 to 12 mW/cm^2^ - Fig.S1 and S2). This is the simplest system to illuminate simple cultures of typically 3 mL, for either overnight cultures or specific, programmed, durations. Specific illumination patterns can also be accommodated by programing the Arduino. The inclination of the tubes and LEDs was designed to increase gas exchange during shaking in the incubator (See Fig.S2 and Supplementary File 1).

#### eVOLVER

The eVOLVER platform, adapted from Wong *et al*. 2018^19^, comprises 16 independent culture units, in which a glass vial can be inserted (Fig.1C), each allowing a single automated yeast culture of 10-25 mL. For each unit, the temperature (heaters and thermistor) and stirring (rotating magnet in vial combined with disc-magnets on a rotating fan) are controlled and the growth rate (infra-red LED and photodiodes) can be monitored semi-quantitatively.

Optogenetics was not included in the original design of the eVOLVER platform, but the addition of a LED in their culture unit PCB to illuminate the culture from the side of the device was anticipated, and we indeed took advantage of this option. We also added a photoresistor so that absorption of blue light in the medium can be monitored. Given that beta-carotene interacts with blue light^30^, its production in cells could be estimated using the blue-LED-photoresistor pair (Supplementary File 2) to obtain qualitative estimations of the cell density and beta-carotene production within the eVOLVER. To further increase the illumination of the cultures, we designed a custom lid: in the hollow lid normally sealed using a removable silicon septum, an autoclavable cytometry tube is placed. Additional LEDs can be stacked inside this tube to illuminate the inside of the medium. Here, we distinguish between the side-LED of the original eVOLVER design (6mW/cm^2^ – external illumination) and the additional LEDs inserted from the top (internal illumination). Adding 1, 2, 3, or 4 additional LEDs was found to correspond to adding 6.7, 8.3, 9.3 and 9.4 mW/cm^2^ of intensity, respectively, in the culture medium.

In our design, each eVOLVER unit is controlled by an individual Arduino Nano and each Arduino is connected to a computer via USB. Compared to the Khalil lab’s fully integrated eVOLVER platform^19^, we propose this more DIY and straightforward design for biologists with minimal knowledge of electronics, since our design circumvents the need to order and assemble various cards and PCBs (see Supplementary File 2). In terms of software, we developed a Node-Red program (detailed in Supplementary File 2 and available on Github) to communicate with the Arduinos in real-time, via the Firmata Arduino Template. This software handles the user interface used to set up and launch experiments, PID used to control temperature, illumination control and data-output. Using this platform containing 16 functional units and an efficient user-interface, we are able to quickly and easily vary multiple parameters at the same time (illumination, volume, stirring, temperature, strain) and thus achieve a high experimental throughput.

#### OptoFlasks

Flasks are the standard culture container used to test strains before scaling up to pilot-scale bioreactors. In order to illuminate flasks, we arranged LEDs into petri-dishes, which were used as illumination stands to illuminate them from the bottom (Fig.1D, S3). LEDs were soldered in a circular pattern such that they line the outermost area of the bottom of the flask. Thus, in a shaking incubator, when the culture medium is driven to the sides of the flask and rotates along the round bottom of the flask, the exposure of the culture to light will be maximized. LEDs are soldered such that each receives 20mA current, resulting in a measured 12mW/cm^2^ intensity per LED, similar to the additional LEDs of eVOLVER and the maximal value of the OptoTubes’ LEDs. Here, we used both flat-bottom flasks and indented-bottom (baffled) 250 mL flasks: the indentation creates turbulences in the liquid flow, which in turn incorporates more air in the culture medium and improves gas transfer. We tested volumes of 25 and 50 mL in 250 mL flat and indented flasks.

### Devices and culture conditions influence optogenetic activation

To investigate how optogenetics translates at different scales, we tested the four optogenetic devices presented above. Each device has its own intrinsic properties: illumination modalities (disposition, number, controllability of the LEDs), geometries (which impact light diffusion and shaking), and range of culture volumes. Illumination, culture volume and shaking can be altered within each device, so that the influence of these factors on optogenetic activation can be identified.

We used the strain OPTO-EXP from the Avalos Lab^9^, a *Saccharomyces cerevisiae* CEN.PK2-1C-derived strain that contains the light-activated EL222-VP16 transcription factor under constitutive expression (pPGK1 promoter). Upon blue light illumination, a change of conformation induced via its flavin chromophore allows EL222 to dimerize, binds to its synthetic cognate pC120 promoter and activate the transcription of the downstream Green Fluorescent Protein (GFP) thanks to its VP16 transactivation domain (Fig.2A).

**Figure 2.**
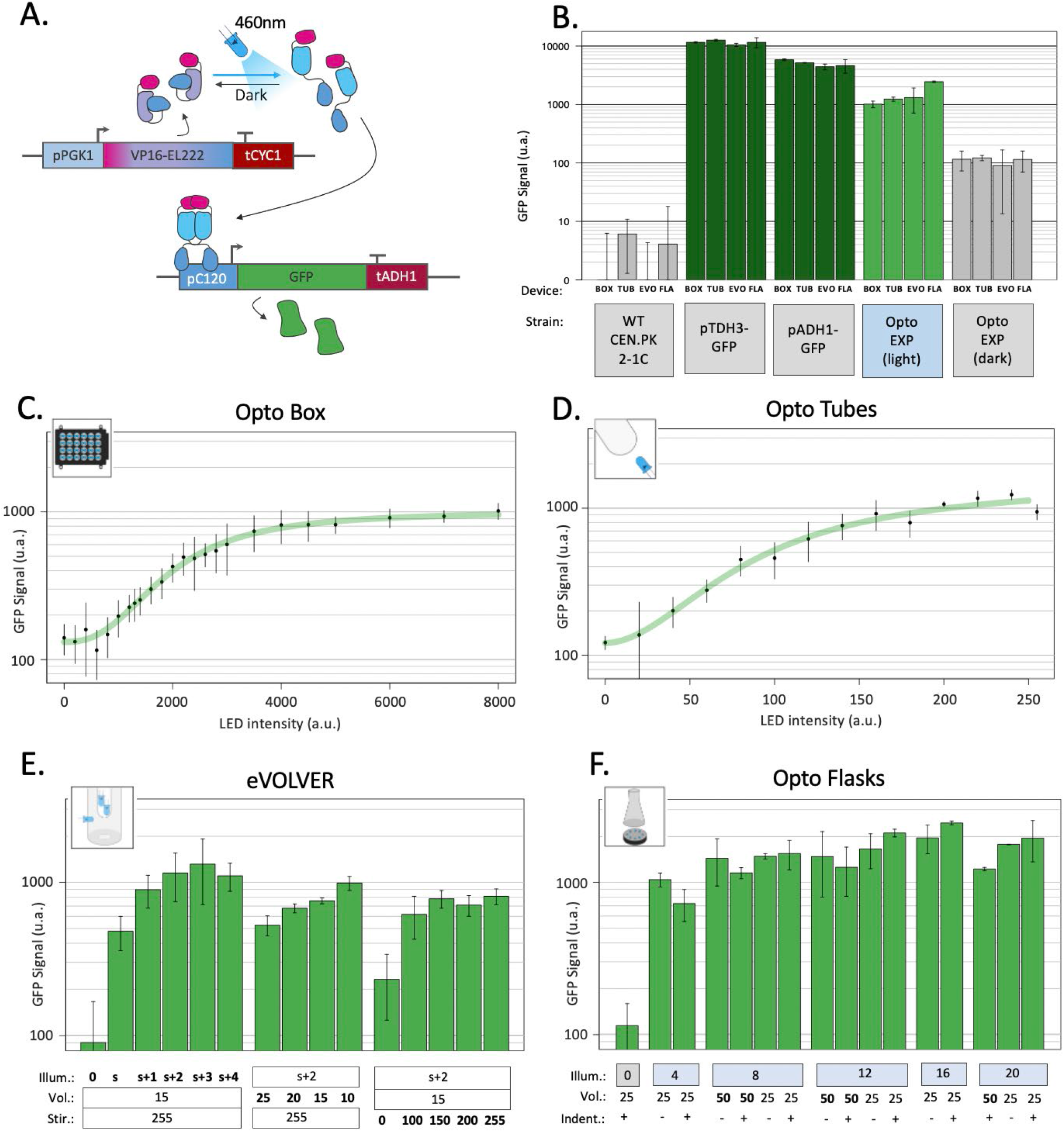
Optogenetic activation in different devices. **(A)** The EL222 optogenetic system responds to blue light which activates the transcription of genes under the control of the pC120 promoter (here a GFP, in the OPTO-EXP strain). Adapted from Zhao *et al*. 2018^9^. **(B)** Fluorescence GFP levels of the background strain CEN.PK2-1C; strains carrying GFP under the strong pTDH3 and medium pADH1 constitutive promoters; and highest levels of fluorescence reached with the OPTO-EXP strain in the light and in the dark in each the four different illumination devices (BOX: OptoBox, TUB: OptoTubes, EVO: eVOLVER, FLA: OptoFlasks). **(C)** OPTO-EXP optogenetic activation in the OptoBox. Cumulative LED intensities from 0 to 8000 u.a. correspond to 0 to 4 mW/cm^2^ per LED. **(D)** OPTO-EXP optogenetic activation in the OptoTubes. LED intensities from 0 to 255 u.a. correspond to 0 to 12 mW/cm^2^ (Fig.S1) **(E)** OPTO-EXP optogenetic activation in the eVOLVER, with variation of illumination (number of LEDs: s+2 corresponds to the side-LED and 2 additional LEDs inserted via the lid), volume (mL) and stirring (u.a., 0 to 255). The side-LED corresponds to 6 mW/cm2, and 1, 2, 3, and 4 additional LEDs add 6.7, 8.3, 9,3 and 9.4 mW/cm^2^ of intensity in the medium. **(F)** OPTO-EXP optogenetic activation in OptoFlasks. Illumination (0, 4, 8, 12, 16 or 20 LEDs on the illumination stand, 12 mW/cm^2^ each). Volume (25 and 50 mL) and the presence of indentation in the 250 mL flask (+ is indented, - is flat) were tested. For all measures, the levels of GFP were determined using cytometry (n > 3); error bars represent the standard deviation.

To test optogenetic activation in different devices and growth conditions, YPD medium was first inoculated with the OPTO-EXP strain at 5.10^6^ cells/mL (OD_600_=0.05), spread across culture containers, placed in illumination devices and illumination started at t0. During growth, as cell density increases, light penetration may be impaired and the amount of light received by each cell may decrease, leading to lower optogenetic activation. Therefore, to determine the best timepoint to obtain a readout of optogenetic activation during growth, we performed a time course experiment, measuring GFP levels every hour using cytometry (Fig. S4 and Methods). We determined that the highest activation per cell was reached at 6 h post-inoculation. Therefore, we standardized the optogenetic experiments using these parameters. As references, we measured fluorescence levels of GFP expressed from the strong and medium constitutive promoters pTDH3 and pADH1 (Fig.2B). To estimate background fluorescence, the autofluorescence of a WT non-producer strain (CEN.PK2-1C) was also measured in the different devices, averaged and subtracted to all datapoints.

By varying the intensity of the LEDs in the Optobox and OptoTubes and measuring the production of GFP (Fig.2C,D), we determined light-response curves. The shaking (250 rpm) and volumes (1 mL and 3 mL) were constant for each device. We obtained two sigmoidal response curves; the dynamic ranges started at 110 arbitrary units of fluorescence (a.u.) and increased to 1014 and 1236 a.u. for the OptoBox and the OptoTubes, respectively, resulting in 9.2- and 11.2-fold, a ∼18% differences in max fluorescence between those two devices. In the OptoBox, maximal activation could almost be reached with one single LED operating at maximal intensity (4000 – 4 mW/cm^2^, for 1 mL). In the OptoTubes, the maximum activation threshold was reached when using maximum intensity (255 – 12 mW/cm^2^, for 3 mL). The effective amount of light reaching the culture volume can be difficult to evaluate due to the geometry of the devices, the type of material and the thicknesses of the materials or media the light has to pass through. However, it is necessary to determine the minimal amount of light needed for maximal optogenetic activation as high doses of blue light can result in phototoxic effects^31,32^.

For eVOLVER and the OptoFlasks we not only varied the illumination but also the culture volume and the agitation properties, *i*.*e*., the stirring speed in eVOLVER and the presence or absence of indentation (*i*.*e*., baffles) in the culture flasks for the OptoFlasks system. Fig. 2E shows that the additional LEDs increase activation compared to the side-LED alone and that the maximal activation is obtained with the side-LED and two to three additional LEDs (s+2 and s+3), reaching about 1190 a.u. Volume also has an impact on optogenetic activation: using a 10 mL culture, activation corresponds to 990 a.u., but only 526 a.u. for 25 mL. Here, reducing the volume 2.5 times yields a 1.9-fold increase in fluorescence. Moreover, in eVOLVER, stirring also proved significant: no stirring leads to an activation of 233 a.u., and low and high stirring (100 to 255 – Arduino PWM values) lead to activation of 730 a.u., corresponding to a 3.1-fold increase. Interestingly, the absence of the stirring appears detrimental, while any other stirring value restores the optogenetic activation level: there, cells don’t simply sit at the bottom of the glass vial.

We reached up to 2453 a.u. in the OptoFlasks (Fig.2F), the highest optogenetic activation of the four devices. Although the impact of the tested volumes (50 and 25 mL) does not appear significant, activation in 25 mL cultures seems to respond more to higher illumination. In general, maximal activation is reached using eight LEDs (12mW/cm^2^ per LED). Using flat or indented flasks did not yield any significant difference in activation.

In general, while the OptoBox, OptoTubes and eVOLVER reach similar maximal activation values of 1015 a.u., activation in OptoFlasks reached an average of 1652 a.u., 1.63-fold higher (Fig.2B), suggesting that the optogenetic activation in the three first devices has potential for improvement. Compared to the basal optogenetic expression in the dark (110 a.u. - leaking), optogenetic activation yielded a 13.7-fold increase in GFP fluorescence. In most of the devices, the optimal activation levels (1505) corresponds to 30% of the pADH1 (5021) and 13% of the pTDH3 promoter activity (11578 a.u.).

### Beta-carotene production is impacted at different scales

Before investigating the efficiency of the optogenetic control of beta-carotene production, we carefully characterized and optimized constitutive beta-carotene production under different culture conditions and across scales. For these experiments, the strain yPH_554, in which the beta-carotene pathway is placed under constitutive expression (*no* optogenetic control), was assayed in the different devices. Culture volume, stirring and light-side-effects have been quantified.

The synthesis of beta-carotene relies on the isoprenoid / mevalonate metabolic pathway. First, acetyl-CoA is channeled towards mevalonate production, then to production of IPP and DMAPP, which are precursors to many other high value-added molecules^33^. Condensation of IPP and DMAPP produces geranyl-pyrophosphate (GPP) and the addition of a second IPP molecule gives farnesyl-pyrophosphate (FPP). FPP can be converted to either farnesol (FOH) via DPP1p; two FPP molecules can condensate to from squalene via ERG9p (the pathway for synthesis of ergosterol and other sterols, essential for cell membranes) or via the addition of another IPP molecule to GGPP, the precursor of the beta-carotene pathway (Fig.3A). To increase GGPP synthesis by yPH_554, the enzymes tHMG1 and CrtE were placed under control of the strong constitutive promoters (pPGK1 and pTDH3 respectively) at the DPP1 locus, while the DPP1 gene was knocked out. Thus, the conversion of acetyl-CoA to mevalonate is increased and the conversion of FPP to FOH is prevented, thus favoring GGPP production. GGPP can then be converted to phytoene (uncolored) by the bi-functional CrtYB enzyme, and subsequently converted to lycopene (red pigment) by CrtI and finally into beta-carotene by CrtYB again. In yPH_554, the genes encoding the CrtYB and CrtI enzymes are inserted at the *ho* locus and both under the strong constitutive promoter pTDH3 (see Methods for more details). This design is based on Rabeharindranto *et al*. 2018^34^, which originated from Verwaal *et al*. 2007^35^.

**Figure 3.**
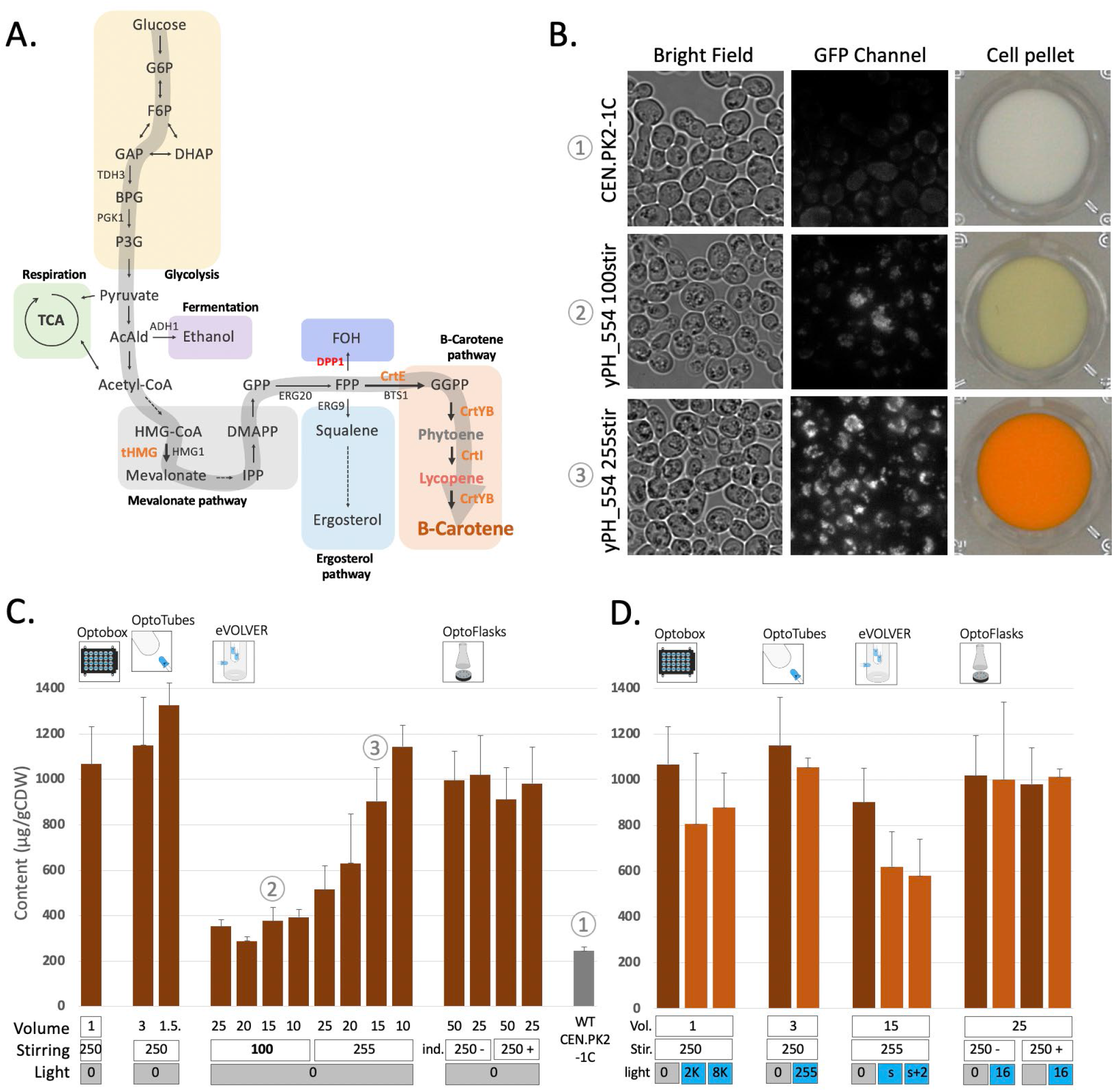
Constitutive beta-carotene production and analysis of the effect of light on beta-carotene accumulation in yeast. **(A)** Beta-carotene pathway. Arrows point to chemical species (see Figure S8 for more details), genes names are indicated in capital letters beside arrows. The large grey arrow represents the carbon flux leading to beta-carotene production. Orange: heterologous genes inserted under constitutive promoters. Red: endogenous gene deletion. **(B)** Microscopic observations and corresponding cell pellets. CEN.PK2-1C (top) strain and constitutive beta-carotene production (yPH_554 – middle and bottom). Growth in YPD at 30 °C for 24 h with low stirring (100 – middle) versus high stirring (255 – bottom). Bright field images and GFP images showing beta-carotene localized in lipid droplets and emitting in the GFP channel (100X objective). See also Fig.S5 **(C)** Constitutive beta-carotene production (content – μg beta-carotene / g cell dry weight) measured after growth in the different devices (see Methods). Volume (mL), stirring/shaking (rpm except for eVOLVER, which is given in arbitrary units, + and - in OptoFlasks represent the presence or absence of indentation, respectively). In this panel, all experiments were performed in the dark (non-optogenetic strain). The WT non-producer strain is CEN.PK2-1C. Sample numbers indicated in encircled numbers in (C) match images in (C). **(D)** Effect of light on constitutive beta-carotene production in the different devices. Dark-orange bars are cultures in the dark, light-orange bars are illuminated cultures. Volume (mL), stirring (rpm or u.a. and indentation presence) are the same as in (C) and light (u.a.) as in Fig.2.

yPH_554 cell pellets appear orange, and cells observed under fluorescence microscopy display numerous fluorescent foci, which are likely to be lipid droplets^36,37^ enriched in beta-carotene (Fig.3B). In fact, these droplets that emit in the GFP channel are orange when observed using a color camera (Fig.S5). A time course of production was carried out to compare production in the different devices: we found that no more beta-carotene was produced after 24 h of culture (Fig.S6). Thus, we standardized our protocol such that cultures are launched at OD_600_ 0.05 and left to grow and produce for 24 h in YPD at 30°C in batches before extraction in dodecane and quantification of beta-carotene via spectrophotometric analysis (based on Reyes & Kao 2018^38^, see Methods).

The effect of culture volume varied across devices (Fig.3C). In the OptoTubes, reducing the culture volume from 3 mL to 1.5 mL yielded a 16% increase in the beta-carotene content, from 1150 and 1330 μg/gCDW (cell dry weight), respectively. Similarly, decreasing the volume in eVOLVER from 25 mL to 10 mL also improved production by 2.2-fold from 515 to 1145 μg/gCDW, respectively. However, this increase was not observed at low stirring, meaning that the limiting factor is probably a combination of low stirring and volume, hence indicating probable differences in aeration between the conditions. At low stirring in eVOLVER (corresponding to the Digital PWM value of 100 sent by the Arduino; the maximum being 255), production is strongly impaired, resulting in an average of 355 μg/gCDW (3.2 times less than the maximum, Fig.3C) and a faintly yellow cell pellet (Fig.3B). In the OptoFlasks, neither the culture volume (50 and 25 mL) nor the presence of indentation in the flasks appeared to affect production. All OptoFlasks conditions yielded slightly less beta-carotene (about 1000 μg/gCDW) than all other devices under their respective optimal conditions.

Beta-carotene is sensitive to blue light: its peak absorption wavelength is about 450 nm^30,39^ and the blue LEDs used to activate the EL222 optogenetic system peak at 461 nm (see Methods). In plants, beta-carotene is even considered to function as a protective molecule against excessive illumination, given its photo-oxidative properties^40^. Therefore, there might be some incompatibility between the blue light-activated optogenetic system and the molecule we wish to produce. Thus, we investigated the impact of blue light on constitutive beta-carotene production (*i*.*e*., *no* optogenetic activation) across devices with the illumination conditions used to activate the optogenetic system. As shown in Fig.3D, in the OptoTubes and OptoFlasks, strong illumination does not impact beta-carotene production. However, strong illumination impacted production in the OptoBox and in eVOLVER, resulting in a drop from 1070 to 845 μg/gCDW (21%) in OptoBox and from 905 to 705 μg/gCDW (22%) in eVOLVER; this effect does not appear to be light dose dependent. It is important to bear these results in mind when interpreting the forthcoming results of the experiments combining optogenetics and beta-carotene production.

In conclusion, we optimized the culture conditions such that the constitutive level of beta-carotene production varied only moderately across devices, ranging from 1000 to 1200 μg/gCDW. The culture conditions were especially improved in the eVOLVER and we showed that culture volume and stirring impacted the production the most. Moreover, beta-carotene production and/or accumulation was only mildly impacted by illumination in OptoBox and eVOLVER.

### Optogenetic control of beta-carotene production in different devices

In the two previous parts, to assess the influence of illumination, volume and stirring across devices, we used two different strains: the OPTO-EXP strain, in which GFP transcription is controlled by the EL222 optogenetic system via its pC120 promoter; and the constitutive beta-carotene producer strain yPH_554, which expresses all four carotenogenic genes under strong constitutive promoters.

Now, we combine these genetic systems into the strain yPH_551, in which the EL222 optogenetic system controls beta-carotene production. tHMG, CrtE, and CrtI were inserted into OPTO-EXP under the control of the constitutive promoters, similarly to the genetic design of yPH_554. However, CrtYB, which recapitulates two enzymatic reactions necessary to produce beta-carotene, was placed under the control of the pC120 optogenetic promoter (see Methods). Thus, in strain yPH_551, the expression of CrtYB acts a light-activated valve for beta-carotene production (Fig.4A).

**Figure 4.**
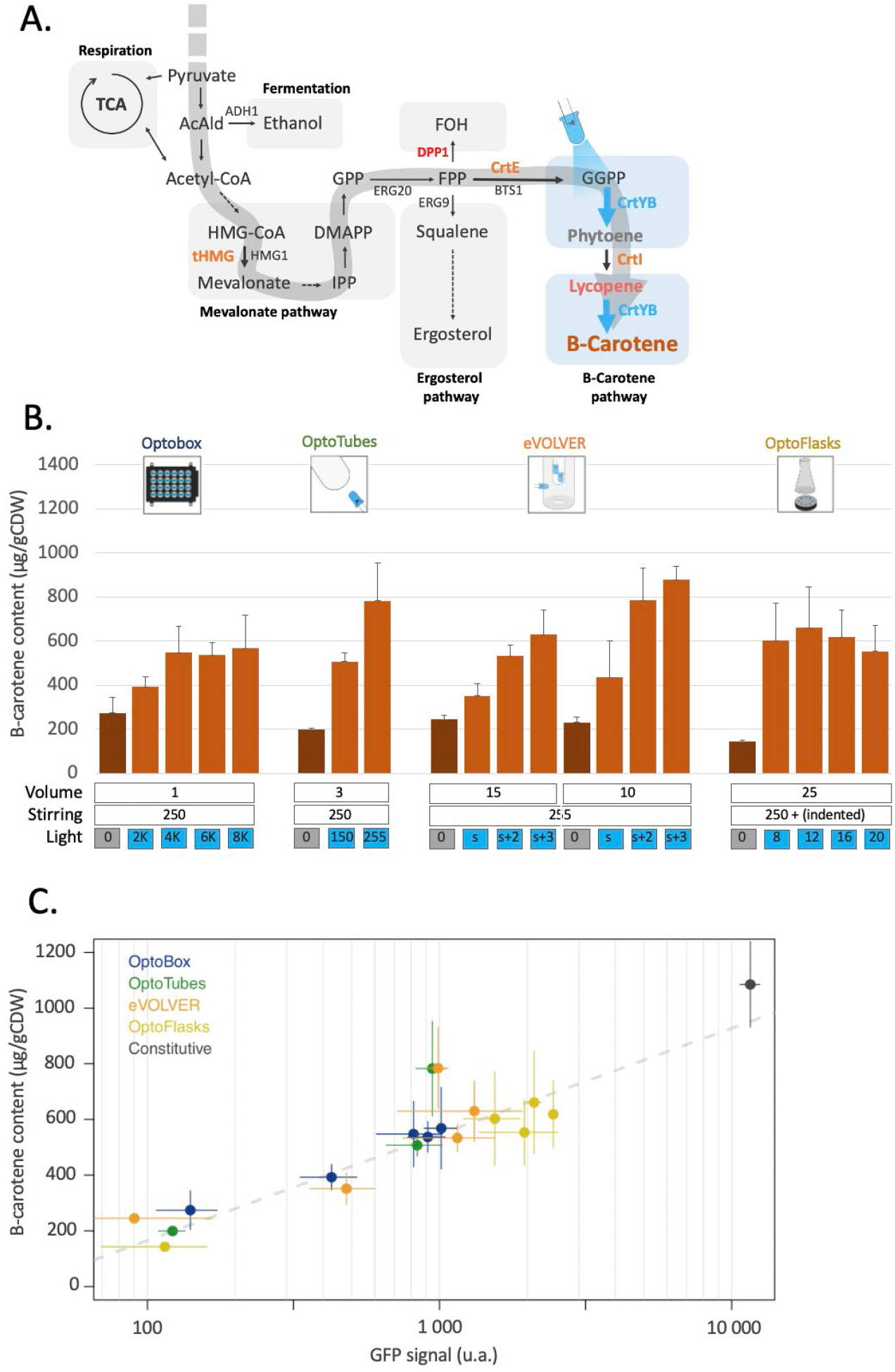
Light-activated beta-carotene production. **(A)** Design leading to light controlled beta-carotene production: in the optogenetic strain, only CrtYB is under the control of the pC120 promoter (yPH_551). Blue arrows represent light-induced enzymatic reactions. Orange: heterologous carotenogenic genes inserted under constitutive promoters. Red: deleted endogenous genes. Blue: optogenetically controlled reaction. **(B)** Beta-carotene production quantification from the different devices. Dark-orange bars correspond to cultures in the dark, light-orange bars correspond to illuminated cultures. Volumes (mL), stirring (rpm except u.a. for eVOLVER) and illumination (u.a.) are indicated (n > 3). **(C)** Light-activated beta-carotene production (Fig.4B) *versus* corresponding optogenetic activation strength (from Fig.2C-F) in the different devices. Grey dashes represent the linear regression fit (adjusted R^2^ = 0.81). The black constitutive point represents the pTDH3 promoter GFP signal and production by the constitutive beta-carotene production strain (yPH_554 - Fig.3C). Error bars are standard deviations.

Fig.4B shows that beta-carotene production by strain yPH_551 was indeed successfully made light-dependent. The average production of 215 μg/g CDW in the dark corresponds to no production of beta-carotene (as confirmed by white cell pellets – see Fig.3C), *i*.*e*., this value is our detection threshold (see Methods).

In the OptoBox and OptoFlasks, optogenetically controlled beta-carotene production saturates at 550 and 610 μg/gCDW respectively. In the OptoTubes and eVOLVER, production increases with the amount of light, to reach a maximum of 780 μg/gCDW in the OptoTubes and 880 in the eVOLVER. The maximal production values in each device were lower than their respective maximal values achieved by constitutive beta-carotene production shown in Fig.3C (52%, 68%, 79%, 62% - for OptoBox, OptoTubes, eVOLVER and OptoFlasks, respectively). Reaching 50 to 80% of constitutive beta-carotene production while optogenetic activation reaches 11% activation of the pTDH3 promoter (Fig.2B) reveals that light induction is efficient, despite the detrimental effects of light on beta-carotene production in the OptoTubes and eVOLVER (Fig.3D)

For most of the conditions tested here, the light-induced production can be related to the previously measured amount of optogenetic activation (Fig.4C), such that conditions yielding more optogenetic activation result in higher beta-carotene production. Here, the production is proportional to the log_10_ of the optogenetic activation value. The consequence of this relationship is that only about 8% of the strength of the pTDH3 promoter is required to reach 50% of the possible beta-carotene production. In other words, even though the Opto-EXP optogenetic system has comparatively low transcriptional strength, in our case, this system produces a good amount of beta-carotene compared to production from a strong constitutive promoter, perhaps because the controlled enzyme, CrtYB, does not conduct the rate-limiting step of the pathway^35^. Finally, the outliers in Fig.4C suggest that culture conditions could probably be further improved to increase production even more.

## Discussion

To test the scalability of the use of optogenetic to control bioproduction and identify its challenges, we built four illumination devices at different lab scales. These devices mainly vary with respect to the culture volumes they hold (from 1 to 50 mL); in their geometry, which impacts shaking and illumination; and in their illumination modalities, which impacts cells exposure to light. Each optogenetic device corresponds to a standard lab-scale culture device, and alternative designs exist for each device and purpose: screen straining in imaging plates like the OptoBox^18,41–43^, illumination of starter cultures in OptoTubes^44^, monitoring multiple culture parameters in minibioreactors such as eVOLVER^19,45,46^, or use of larger culture volumes for various purposes in OptoFlasks^47^. Measuring the intensities of the LEDs allows us to compare the relative amounts of input light across these devices. However, due to the unique geometry of each device (position of the LEDs, light absorption, diffraction, and reflection) and their different culture volumes (light penetration), it remains difficult to precisely estimate the actual quantity of light received by each cell; thus, testing the activation of a reporter gene like GFP remains the best way to take these technical aspects into consideration in combination with potential biological limitations.

We concluded that optogenetic activation reached an average maximum of 13% of the pTDH3 promoter across devices. Estimations of the quantity of light per milliliter of culture revealed that between 1 and 4 mW/cm^2^/mL was sufficient for maximal activation in the different devices. The variations across devices indicate that other factors impact optogenetic activation in addition to the amount of light, even though measuring the effective amount of light entering and staying in the media remains difficult. Light distribution has been actively studied using single-particle and fluid dynamics models to optimize illumination of photobioreactors (mostly cultures of photosynthetic microorganisms) and predict the amount of light received by each cell over time^48,49^; and microbial optogenetics could indeed profit from such approaches.

In eVOLVER, the volume and stirring parameters strongly impacted optogenetic activation. Indeed, reducing the volume from 25 to 10 mL with maximum stirring and s+2 illumination improved activation by almost 2-fold. Furthermore, a lack of stirring resulted in a 3-fold decrease in activation, while it remained constant between low (100) and strong (255) stirring. One can interpret these results from two points of view: in terms of gas exchange in the medium and in terms of light availability. On one hand, a decrease in volume will increase the surface-to-volume ratio in the vial and improve gas exchange, and more specifically the oxygen transfer rate (OTR), a crucial element taken into consideration in bioreactors. Similarly, increasing the stirring will create a bigger vortex, resulting in comparable effects. Since only the absence of stirring (and not an increase in stirring) impacts activation, we suggest that the limitation of the OTR may be relatively low in this case, justified by the fact that the cultures measured in our experiments were only 6 h old. On the other hand, these effects may relate to the amount of light received per cell. For a set amount of light, a lower culture volume (lower cell number) could result in a higher amount of light per cell. Moreover, in eVOLVER, a high volume (25 mL) implies that some parts of the culture volume (and therefore, cells) are lying above the LEDs (*i*.*e*., the poorly illuminated area - Fig.1C) and a higher volume also leads to lower light penetration; both of these factors result in a decrease in the amount of light per milliliter. In addition, in the absence of stirring, the cells lie still on the bottom of the glass vial and remain far from the light sources due to the position of the LEDs, which are placed at the top and on the side in eVOLVER. These results emphasize the importance of the illumination design within each of the different devices, which may not necessarily be well-suited for larger culture volumes.

Using our genetic design, constitutive beta-carotene production achieved a content of 1000 to 1300 mg/gCDW in the different devices. Beta-carotene production was sensitive to the culture volume and stirring. Reducing stirring in eVOLVER resulted in a 3-fold drop in production and increasing the volume to 25 mL led to a 2-fold drop (compared to production at 255 stirring of 10 mL). Culture volume also had an effect in the OptoTubes, but not in the OptoFlasks. This highlights the need to optimize the culture conditions of every device to obtain a functional strain in each and across scales. Since both higher stirring and lower volume impact beta-carotene production, we suggest that improved gas-exchange favors beta-carotene production by impacting cellular metabolism. At a sufficient concentration of oxygen (which is constrained by gas-exchange), *S. cerevisiae*, a Crabtree-positive species, undergoes both fermentation and also respiration^50^, which consequently increases the acetyl-CoA pool directed to the Krebs cycle as well as other precursors to beta-carotene.

The effects of volume and stirring on our two different genetic components were found to overlap: more stirring and a lower volume improved both optogenetic activation (for optimal illumination reasons) and beta-carotene production (for metabolic reasons), making those two genetic components easily compatible. This compatibility may not always be consistent for every system; hence, it is crucial to anticipate and evaluate these effects independently before combining optogenetic control with the pathway of interest.

We saw that the constraints of the optogenetic system and the beta-carotene pathway in each device are recapitulated in the optogenetic beta-carotene strain: in general, more light resulted in more production, and a smaller volume in eVOLVER favored the light-activated production. Production levels reached 50 to 80% of those from constitutive production. Since we saw that production was proportional to the log_10_ of the expression of the enzyme, this lower production reached using optogenetics is likely due to the relatively low transcriptional strength of the pC120 optogenetic promoter. This could be solved by adding more copies of pC120-CrtYB or by using a different^9^ or more recent and stronger optogenetic system^51–53^.

Although we found that more activation (optogenetic or constitutive) yielded more beta-carotene production, this straightforward relationship may not always stand^54,55^. Indeed, relative enzymatic levels can have an impact on the production of different metabolites (and this has even been demonstrated for beta-carotene production^34,56^). Therefore, fine-tuning the optimal enzyme concentration with optogenetics could result in higher yields in different designs and for different metabolic pathways. More importantly, if higher expression yields more production, it can also lead to a higher burden and impact growth and strain stability, and therefore impact the production titers. Using optogenetics could enable precise control of the levels of enzymes using dynamic control based on feedback from the expression levels and growth rate^57^.

Optogenetics also allows for precise control of gene expression in terms of timing. Here, we activated the cells constantly during a 24 h culture. Nevertheless, other studies have indicated that starting the illumination at later phases of growth or using pulsed patterns of illumination can improve production: Zhao *et al*. 2021^51^ distinguished the growth, induction and production phases and differently optimized the illumination patterns to produce specific chemicals and Raghavan *et al*. 2020^12^ defined the optimal illumination onsets to start the production.

This question of timing can be especially important when the production of the chemical of interest creates a burden on the cell and therefore slows down growth and/or creates a stress. In our study, beta-carotene production did not appear to produce a burden (Fig.S7); however, if this were the case, the ease of timing optimization and control using optogenetics could allow for improved titers (g/L of culture) and yields (g/g of carbon source), not only content (g/gCDW - shown in this study). To further reduce this burden, dynamic control of bioproduction could be employed to let the cells recover from the production stress and then resume production with a renewed high productivity^57^.

The four optogenetic devices presented here are lab-scale culture devices. Scaling-up optogenetics to industrial culture volumes will generate different constraints on optogenetic activation and yeast metabolism, although bioreactors often have better control over culture parameters than lab-scale devices (pH, dissolved O_2_ and CO_2_, temperature). Other papers have already demonstrated optogenetic control of bioproduction in up to 5 L bioreactors^51^ and we look forward to seeing how optogenetics can be scaled-up to industrial settings^57^.

## CONCLUSION

We suggest a three-step approach to apply optogenetics to bioproduction control: characterize and optimize each of the optogenetic and the bioproduction components independently, before combining them. Indeed, optogenetic activation can vary across devices and the production behavior of a strain can be significantly affected by seemingly minor differences in culture conditions. Overall, this approach can help to reveal incompatibilities between the genetic components and eliminate optogenetic-dependent or pathway-dependent confounding factors.

While optogenetic systems are becoming better and illumination devices more widespread, the use of optogenetics to control microbial systems and its application to bioproduction is still a biotechnology in its infancy. Scalability remains a technical challenge and this study sets the stage for every step of the strain development process at the lab scale. Coming up next will be small and larger bioreactors before industrial scales. The promises of optogenetics for bioproduction lie in the controllability it offers to achieve more robust bioprocesses and apply real-time dynamic control of growth *versus* production, wherein lies an important resource allocation trade-off, with the overall goal of improving yields and making bioproduction an economically viable technology for sustainable production of any type of chemical or protein for multiple industrial sectors.

## MATERIALS AND METHODS

### Construction of Yeast Strains

The GFP optogenetic strain OPTO-EXP originated from Zhao *et al*. 2018^9^. Using the CEN.PK2-1C background, pPGK1-EL222 (constitutive expression of the light-activated transcription factor) and pC120-GFP (optogenetic promoter to which EL222 binds to activate the GFP transcription) were inserted at the *Δhis3* locus, with a functional HIS3 copy (*Cg*HIS3).

For constitutive beta-carotene production, constructs were made using Rabeharindranto *et al*. 2019^34^’s DNA material in which a trifusion enzyme CrtYBekI was developed. In plasmid pHR0016, the pGal1-10 promoter was replaced with pTDH3 using the Gibson assembly^58^, to obtain plasmid pPH_386. Similarly, the bidirectional pGAL1-10 controlling the expression of CrtE and tHMG1 in plasmid pMRI34 was replaced with pPGK1-pTDH3 to build plasmid pPH_350.

In strain CEN.PK2-1C, the trifusion CrtYBekI was inserted at the *ho* locus using CRISPR^59^. In brief, a gRNA targeting the *ho* locus was designed and inserted into the pML104 plasmid containing cas9, the gRNA expression cassette and the URA3 marker (pML04-ho, built using restriction enzyme digestion and ligation according to Laughery *et al*. 2015^59^). The template DNA strand was amplified from the plasmid pPH_386, containing pTDH3-CrtYBekI and two homologous arms of 90 bp targeting the *ho* locus. Upon transformation using the LiOAc Gietz method^60^, both the pML104-ho plasmid and the template strand were added to the mix. In brief, Cas9 will cut repeatedly at the *ho* locus and favors homologous recombination with the template DNA at this locus, which acts as selective pressure against perfect double stranded break repair. After transformation, cells were plated on Complete Synthetic Medium (CSM)-URA. Clones were isolated and screened using OneTaq DNA polymerase (NEB). The pML104-ho plasmid was then cured using 5-fluoroorotic acid (5-FOA) CSM. The trifusion design^34^ was broken down to the natural bifusion CrtYB and CrtI using the same CRISPR method (pML104-ek, and template from YIplac211), resulting in pTDH3-CrtYB-pTDH3-CrtI at the *ho* locus. In addition, using a gRNA targeting the *DPP1* locus (pML104-DPP1), CrtE and tHMG were inserted under the constitutive pPGK1 and pTDH3 promoters (pPH_350), both of which favor carbon flux to increase production of the GGPP precursor (at the same time, DPP1 was deleted, which was shown to improve production^34^).

To achieve optogenetic control of beta-carotene production (yPH_551), the background strain was OPTO-EXP-mCherry (yPH_463), in which the GFP reporter of the optoEXP system at the *HIS* locus of the OPTO-EXP strain was replaced with mCherry (using pML104-GFP) to prevent spectral overlap in microscopy (beta-carotene localizes in lipid droplets that are visible in the GFP channel and would therefore overlap with the GFP signal). The CrtYBekI trifusion was placed under the control of the pC120 optogenetic promoter in plasmid pHR0016 (amplified from the plasmid EZ-L83) using the Gibson assembly^58^. Then, the CrtYBekI trifusion was inserted at the *ho* locus in strain OPTO-EXP-mCherry (yPH_463) and broken down using CRISPR, similarly to the constitutive yPH_554 strain. In addition, as for yPH_554, CrtE-pTDH3-pPGK1-tHMG was inserted at the *dpp1* locus.

Plasmids used in this study

| Plasmid | Derived from | Description | Source |
| --- | --- | --- | --- |
| **pHR0016** | - | pGAL1-10-CrtYB(ek)I | Rabeharindranto 2019<sup>34</sup> |
| **pPH_386** | pHR0016 | pTDH3-CrtYB(ek)I | This study |
| **pPH_371** | pHR0016 | pC120-CrtYB(ek)I | This study |
| **pMRI34** | - | CrtE-pGal1-10-tHMG1 | Rabeharindranto 2019<sup>34</sup> |
| **pPH_350** | pMRI34 | CrtE-pTDH3-pPGK1-tHMG1 | This study |
| **YIplac211** | - | pTDH3-CrtYB, pTDH3-CrtI, pTDH3-CrtE | Verwaal 2007<sup>35</sup> |
| **EZ-L83** | - | pC120-GFP | Zhao 2018<sup>9</sup> |
| **pML104** | - | Cas9, gRNA scaffold (URA3) | Laughery 2015<sup>59</sup> |
| **pML104-ho** | pML104 | gRNA targeting *ho* promoter | This study |
| **pML104-DPP1** | pML104 | gRNA targeting DPP1 CDS | This study |
| **pML104-GFP** | pML104 | gRNA targeting GFP CDS | This study |
| **pML104-ek** | pML104 | gRNA targeting the (ek) linker | This study |

Strains used in this study

| Yeast strain | Genotype | Source |
| --- | --- | --- |
| **CEN.PK2-1C** | *MATa; ura3-52; trp1-289, leu2-3_112; his3D1;; MAL2-8c; SUC2* | Euroscarf (ref 30000A) |
| **yPH_545** | *CEN.PK2-1C*<br>*ho::pTDH3-GFP* | This study |
| **yPH_550** | *CEN.PK2-1C*<br>*ho::pADH1-GFP* | This study |
| **OPTO-EXP**<br>(YEZ139) | *CEN.PK2-1C*<br>*his3D1::pPGK1-EL222-pC120-GFP (CgHIS3)* | Zhao 2018<sup>9</sup> |
| **yPH_463**<br>(OPTO-EXP-mCherry) | *CEN.PK2-1C*<br>*his3D1::pPGK1-EL222-pC120-mCherry (CgHIS3)* | This study |
| **yPH_554** | *CEN.PK2-1C*<br>*Δho::pTDH3-CrtYB-pTDH3-CrtI* | This study |
| | $\Delta dpp1::CrtE$-pTDH3-pPGK1-tHMG | |
| yPH_551 | OPTO-EXP-mCherry (yPH_463)<br>$\Delta ho::pC120$-CrtYB-pTDH3-CrtI<br>$\Delta dpp1::CrtE$-pTDH3-pPGK1-tHMG | This study |

### Growth Conditions

All strains were grown in standard, home-made, filter-sterilized YPD [yeast extract 1% *w/v* (BD 212750), peptone 2% *w/v* (BD 211677) and D-glucose 2% *w/v*] at 30 °C. All devices except for eVOLVER, were placed in incubators with shaking at 250 rpm and protected from ambient light. For strain selection after transformation, CSM-URA was made with Yeast Nitrogen Base (YNB) without amino acids 0.67% *w/v* (BD 291940), D-glucose 2% *w/v*, CSM dropout-URA 0.08% (MP Bio 114500022). To prepare 5-FOA medium, 50 mL/L of 1g/L uracil solution and 0.8 g/L 5-FOA were added to CSM-URA.

### Illumination devices

#### LED intensity

Wurth Elektronik 151054BS04500 LEDs with a peak wavelength of 461 nm were used for all devices. Intensity measurements were conducted using a power meter (TOR Labs S120C) and divided by the sensor surface (0.7088 cm^2^). The intensities of the LEDs depends on their connection. Optobox LEDs have a maximum intensity of 4 mW/cm^2^ due to the resistance present in the PCB design. Similarly, in eVOLVER, the side-LED intensity is limited to 6 mW/cm^2^ by the 82 Ohm resistance. The additional LEDs in the eVOLVER and the OptoTubes LEDs are directly connected to the digital pins of an Arduino, such that their current is only dependent on the Arduino, reaching about 20 mAmp and yielding 12 mW/cm^2^ intensity. In the OptoFlasks, the illumination stand was designed with the 20 mAmp constraint mentioned earlier; four LEDs in series was computed to reach a similar current per LED, such that their intensity was also 12 mW/cm^2^.

#### OptoBox

Based on Gerhardt *et al*. 2016^18^, the OptoBox (*a*.*k*.*a*. the Light Plate Apparatus) was assembled using ordered PCBs, soldered LED-sockets, inserted blue LEDs and 3D-printed cases. LED calibration was performed using a power meter (TOR Labs S120C) and increasing illumination steps to achieve homogeneous illumination between all LEDs and wells at the same illumination intensity. Black 24-well cell imaging plates (Eppendorf 0030741005) were used to prevent light leaking between wells and sealed with aluminum foil to prevent light leakage and culture spillage and evaporation. A 25-μm film lies at bottom of the well plate to allow high gas permeability and UV-light transparency, according to the manufacturer. For an experiment, 1 mL of inoculated medium is placed into each well. A sticking aluminum sheet is used to seal the wells, and the lid is placed in top. The imaging plate is placed in the previously programmed (via its SD card) OptoBox, which is then connected to a 5 V power supply. The device sits in a 30 °C incubator shaking at 250 rpm.

#### OptoTubes

LEDs are encased in a box at regular intervals (4 × 2 LEDs) and the tubes are held on top, with their bottom almost touching the LED, via a 3D printed holder (Fig.S2 and Supplementary File 1). LEDs are directly connected to an Arduino PWM pin and were measured to result in a LED intensity of 12 mW/cm^2^. Pictures and schemes are shown in Fig.S2. To run an experiment, 3 mL (generally) of inoculated medium is placed into a 14 mL glass test tube. The tubes are closed with a cellulose cap, placed in the OptoTubes device, and the Arduino (previously programmed with the Arduino software for the chosen light intensities) is connected to 12 V power supply. The device also sits in a 30 °C incubator shaking at 250 rpm.

#### eVOLVER

eVOLVER was built following the instructions of Wong *et al*. 2018^19^. To launch an experiment, autoclaved vials are filled with inoculated medium and closed with a simple lid or with the custom-lid (briefly detailed below and more extensively described in Supplementary File 2). The lid is loosely placed on top of the vial, *not* tightly screwed on, to allow for gas exchange. The vials are then placed in the aluminum sleeves (which are used for temperature control) of each eVOLVER unit. There, the additional LEDs, are placed and stacked by hand into each custom-lid. Using Node-RED software on the user interface, the culture parameters can be set (stirring, illumination from the side-LED, temperature), measure cycles modified, and names and notes added for each unit independently.

The lid of the glass vial used in eVOLVER is a hollow cap, normally closed with a removable silicone septum. To improve illumination, the cap was modified to accommodate insertion of LEDs that reach into the medium (custom-lid). For this, autoclavable cytometry tubes (5 mL Polypropylene Round-Bottom Tubes; BD Falcon REF 352063) are held tightly inside the cap hole using a larger piece of silicon (autoclavable) tubing. Using this design, additional LEDs can be placed inside the culture vial without getting wet and while maintaining sterility (Supplementary File 2). In the intensity measurements, we distinguished the side-LED of the original eVOLVER design (6 mW/cm^2^ - a 82 Ohm resistor sets its intensity^19^) from the additional LEDs inserted from the top (12 mW/cm^2^ each, independently). However, combining several additional LEDs in the custom-lid does not result in a linear increase in total intensity. Since the LEDs are stacked on top of each other in a plastic (quite opaque) cytometer tube, adding 1, 2, 3, or 4 additional LEDs was measured to correspond to adding 6.7, 8.3, 9,3 and 9.4 mW/cm^2^ of intensity, respectively, (measured at the tip of the tube – Fig.S1) in the culture medium.

#### OptoFlask

The OptoFlask illumination stand is made by connecting standard LEDs in series of four in order to have 20 mA per LEDs (similar to the Arduino output, and recommended in the LED specs datasheet) with a 12 V input (DC power connector socket). Several series of four LEDs can be connected in derivation such that all LEDs remain at 20 mA per LED, resulting in a proportional increase in light intensity in the different illumination stands containing 4, 8, 12, 16 or 20 LEDs. To set up an experiment, 250 mL flasks (flat or indented -baffled – bottom) are filled with inoculated media and sealed with a cotton-cap, held with a thin rubber-band. The 250 mL flasks are positioned on the illumination stand, then placed in an oversized metal flask holder in a 30 °C shaking incubator (250 rpm) and held in place with a metal string and lab tape. Then, the stand is connected to the 12 V power supply. See Fig.S3.

### Quantification of optogenetic activation (cytometry)

To quantify the activation of the optogenetic system in the different devices under different growth and illumination conditions, the YEZ139 strain (OPTO-EXP^9^) was streaked onto YPD from a glycerol stock and incubated at room temperature for 48 h. For the preculture, a single colony was picked from the plate and incubated overnight in the dark. The next morning, an aliquot of the preculture was inoculated into YPD to obtain a OD_600_ of 0.05 (5.10^6^ cells/mL) and then dispensed into the different containers for the illuminated cultures. The cultures and illumination were set for 6 hours: 2 Optoboxes, 8 OptoTubes, 5 Flasks, and 16 eVOLVER units can be tested in parallel. After culture, 200 μL of cultures were diluted in 200 μL of PBS and the levels of GFP resulting from the optogenetic activation were quantified using a BD LSR II flow cytometer (BD Biosciences) at an excitation wavelength of 488 nm and emission wavelength of 530 nm. The acquisition settings (voltage) for fluorescence quantification were identical for all experiments. Data were collected for 10,000 cells in each culture and analysis was performed in R using the FlowCore^61^ package.

### Beta-carotene extraction and content estimation

Beta-carotene quantification was adapted from Reyes & Kao 2018^38^. In brief, after inoculation of the media at OD_600_ 0.05 and growth for 24 h, each yeast culture sample was diluted 1:100, the OD_600_ was read and Cell Dry Weight was estimated accordingly using a calibration curve (of equivalences between spectrophotometer and actual freeze-dried weighed samples). Then, 1 mL of culture was transferred to a collection tube (MP Bio). The cells were collected by centrifugation at 11 000 x *g* for 2 min, the supernatant was discarded, 250 μL of acid-washed glass beads (Sigma 425-600um – G8772) and 1 mL of dodecane were added to the cell pellet and the yeast cells were lysed using a FastPrep bead-beater (MP Bio) for five times for 1 min to ensure maximal carotenoid recovery. After cell disruption, the samples were centrifuged (11 000 x *g*, 2 min) to separate cell debris and glass beads, 200 μL of the supernatant was transferred to a 96 well plate and scanned from OD_200_ to OD_700_ using a Spark TECAN analyzer. The A_454_ of the scan was used to estimate the beta-carotene of the dodecane solution using calibration curve prepared with hexane and beta-carotene’s extinction coefficient. By comparing the determined beta-carotene concentration in the dodecane solution with the actual OD and culture volume, the content, yield and titer can be calculated. Note that *non*-beta-carotene-producing cells yield a production value of about 200 μg/g CDW, even though the colonies will appear white, *i*.*e*., this value is the limit of detection.

To view the color of the cell pellets, cultures were concentrated to OD 50 by centrifugation, 200 μL was poured into a 96-well-plate, allowed to sit for 15 minutes, then the plate was color-scanned using a desktop scanner.

### Microscopy

Bright field (BF) and fluorescence images were acquired with MetaMorph software from an Olympus IX83 inverted epifluorescence microscope paired with an Andor CMOS Zyla camera using an 100x objective. Samples were illuminated using a CoolLED pE4000 fluorescence lamp. An exposure time of 200 ms was used to acquire BF images. Images of GFP fluorescence were acquired using excitation and emission wavelengths of 470 and 525 nm, respectively, and an exposure time of 250 ms at 25% light intensity. To acquire images, a small volumes of culture were centrifuged and the cells were simply loaded into a custom-made PDMS chip^62^ to obtain a monolayer of cells. The chip was then placed under the 100x objective for image acquisition.

## DATA AVAILABILITY STATEMENT

The datasets presented in this study can be found in the following Zenodo repository DOI: 10.5281/zenodo.7265419. Code used to analyze the data are available on GitHub at https://github.com/Lab513/DIY_Optogenetics

## AUTHOR CONTRIBUTIONS

SP performed all experiments and participated in all aspects of this work. JRC contributed to the experiments related to beta-carotene production and extraction. MLB, CC, AB, SB, and SC contributed the experiments and associated with the setup and control of the optogenetic devices. SP, BS, TL, GT and PH designed the study and wrote the article.

## ACKNOWLEDGMENTS

We would like to thank Doctor Jose Avalos for kindly providing us with the optogenetic strains and genetic materials. We also thank Guillermo Nevot, William Bretts and the mechanical workshop of the Matter and Complex Systems Lab, especially Oune-Saysavanh Souramasing, for their help with setting up the eVOLVER hardware.

## SUPPLEMENTARY MATERIAL

The Supplementary Material for this article contains 3 supplementary files and 8 supplementary figures.

## REFERENCES

1. Srinivasan P, Smolke CD. Biosynthesis of medicinal tropane alkaloids in yeast. Nature. 2020;585(7826):614–619. doi:10.1038/s41586-020-2650-9

2. Galanie S, Thodey K, Trenchard IJ, Filsinger Interrante M, Smolke CD. Complete biosynthesis of opioids in yeast. Science (80-). 2015;349(6252):1095–1100. doi:10.1126/science.aac9373

3. Chen R, Yang S, Zhang L, Zhou YJ. Advanced Strategies for Production of Natural Products in Yeast. iScience. 2020;23(3):100879. doi:10.1016/j.isci.2020.100879

4. Oud B, Van Maris AJA, Daran JM, Pronk JT. Genome-wide analytical approaches for reverse metabolic engineering of industrially relevant phenotypes in yeast. FEMS Yeast Res. 2012;12(2):183–196. doi:10.1111/j.1567-1364.2011.00776.x

5. Pereira F, Lopes H, Maia P, et al. Model-guided development of an evolutionarily stable yeast chassis. Mol Syst Biol. 2021;17(7):1–18. doi:10.15252/msb.202110253

6. Subramanian I, Verma S, Kumar S, Jere A, Anamika K. Multi-omics Data Integration, Interpretation, and Its Application. Bioinform Biol Insights. 2020;14:7–9. doi:10.1177/1177932219899051

7. Sahu A, Blätke MA, Szymański JJ, Töpfer N. Advances in flux balance analysis by integrating machine learning and mechanism-based models. Comput Struct Biotechnol J. 2021;19:4626–4640. doi:10.1016/j.csbj.2021.08.004

8. Martin JF, Demain AL. Control of antibiotic biosynthesis. Microbiol Rev. 1980;44(2):230–251. doi:10.1128/mmbr.44.2.230-251.1980

9. Zhao EM, Zhang Y, Mehl J, et al. Optogenetic regulation of engineered cellular metabolism for microbial chemical production. Nature. 2018;555(7698):683–687. doi:10.1038/nature26141

10. Lalwani MA, Zhao EM, Wegner SA, Avalos JL. The Neurospora crassa Inducible Q System Enables Simultaneous Optogenetic Amplification and Inversion in Saccharomyces cerevisiae for Bidirectional Control of Gene Expression. ACS Synth Biol. 2021;10(8):2060–2075. doi:10.1021/acssynbio.1c00229

11. Tan SZ, Prather KL. Dynamic pathway regulation: recent advances and methods of construction. Curr Opin Chem Biol. 2017;41:28–35. doi:10.1016/j.cbpa.2017.10.004

12. Raghavan AR, Salim K, Yadav VG. Optogenetic Control of Heterologous Metabolism in E. coli. ACS Synth Biol. 2020;9(9):2291–2300. doi:10.1021/acssynbio.9b00454

13. Lalwani MA, Ip SS, Carrasco-López C, et al. Optogenetic control of the lac operon for bacterial chemical and protein production. Nat Chem Biol. Published online 2020. doi:10.1038/s41589-020-0639-1

14. Senoo S, Tandar ST, Kitamura S, Toya Y, Shimizu H. Light-inducible flux control of triosephosphate isomerase on glycolysis in Escherichia coli. Biotechnol Bioeng. 2019;116(12):3292–3300. doi:10.1002/bit.27148

15. Ding Q, Ma D, Liu GQ, et al. Light-powered Escherichia coli cell division for chemical production. Nat Commun. 2020;11(1):1–14. doi:10.1038/s41467-020-16154-3

16. Zhao EM, Suek N, Wilson MZ, et al. Light-based control of metabolic flux through assembly of synthetic organelles. Nat Chem Biol. 2019;15(6):589–597. doi:10.1038/s41589-019-0284-8

17. Lalwani MA, Kawabe H, Mays RL, Hoffman SM, Avalos JL. Optogenetic Control of Microbial Consortia Populations for Chemical Production. ACS Synth Biol. 2021;10(8):2015–2029. doi:10.1021/acssynbio.1c00182

18. Gerhardt KP, Olson EJ, Castillo-Hair SM, et al. An open-hardware platform for optogenetics and photobiology. Sci Rep. 2016;6(June):1–13. doi:10.1038/srep35363

19. Wong BG, Mancuso CP, Kiriakov S, Bashor CJ, Khalil AS. Precise, automated control of conditions for high-throughput growth of yeast and bacteria with eVOLVER. Nat Publ Gr. 2018;36(April). doi:10.1038/nbt.4151

20. Motta-Mena LB, Reade A, Mallory MJ, et al. An optogenetic gene expression system with rapid activation and deactivation kinetics. Nat Chem Biol. 2014;10(3):196–202. doi:10.1038/nchembio.1430

21. Bogacz-Radomska L, Harasym J. β-Carotene-properties and production methods. Food Qual Saf. 2018;2(2):69–74. doi:10.1093/fqsafe/fyy004

22. Ribeiro BD, Barreto DW, Coelho MAZ. Technological Aspects of β-Carotene Production. Food Bioprocess Technol. 2011;4(5):693–701. doi:10.1007/s11947-011-0545-3

23. Paddon CJ, Westfall PJ, Pitera DJ, et al. High-level semi-synthetic production of the potent antimalarial artemisinin. Nature. 2013;496(7446):528–532. doi:10.1038/nature12051

24. Nowrouzi B, Li RA, Walls LE, et al. Enhanced production of taxadiene in Saccharomyces cerevisiae. Microb Cell Fact. 2020;19(1):1–12. doi:10.1186/s12934-020-01458-2

25. Hansen NL, Miettinen K, Zhao Y, et al. Integrating pathway elucidation with yeast engineering to produce polpunonic acid the precursor of the anti-obesity agent celastrol. Microb Cell Fact. 2020;19(1):1–17. doi:10.1186/s12934-020-1284-9

26. Benzinger D, Ovinnikov S, Khammash M. Synthetic gene networks recapitulate dynamic signal decoding and differential gene expression. Cell Syst. 2022;13(5):353-364.e6. doi:10.1016/j.cels.2022.02.004

27. Shaw WM, Yamauchi H, Mead J, et al. Engineering a Model Cell for Rational Tuning of GPCR Signaling. Cell. 2019;177(3):782-796.e27. doi:10.1016/j.cell.2019.02.023

28. Liu R, Liu L, Li X, Liu D, Yuan Y. Engineering yeast artificial core promoter with designated base motifs. Microb Cell Fact. 2020;19(1):1–9. doi:10.1186/s12934-020-01305-4

29. Duplus-Bottin H, Spichty M, Triqueneaux G, et al. A single-chain and fast-responding light-inducible cre recombinase as a novel optogenetic switch. Elife. 2021;10:1–52. doi:10.7554/eLife.61268

30. Pénicaud C, Achir N, Dhuique-Mayer C, Dornier M, Bohuon P. Degradation of β-carotene during fruit and vegetable processing or storage: Reaction mechanisms and kinetic aspects: A review. Fruits. 2011;66(6):417–440. doi:10.1051/fruits/2011058

31. Enwemeka CS, Baker TL, Bumah V V. The role of UV and blue light in photo-eradication of microorganisms. J Photochem Photobiol. 2021;8:100064. doi:10.1016/j.jpap.2021.100064

32. Grangeteau C, Lepinois F, Winckler P, Perrier-Cornet JM, Dupont S, Beney L. Cell death mechanisms induced by photo-oxidation studied at the cell scale in the yeast Saccharomyces cerevisiae. Front Microbiol. 2018;9(NOV):1–8. doi:10.3389/fmicb.2018.02640

33. Clomburg JM, Qian S, Tan Z, Cheong S, Gonzalez R. The isoprenoid alcohol pathway, a synthetic route for isoprenoid biosynthesis. Proc Natl Acad Sci U S A. 2019;116(26):12810–12815. doi:10.1073/pnas.1821004116

34. Rabeharindranto H, Castaño-Cerezo S, Lautier T, et al. Enzyme-fusion strategies for redirecting and improving carotenoid synthesis in S. cerevisiae. Metab Eng Commun. 2019;8(December 2018):1–11. doi:10.1016/j.mec.2019.e00086

35. Verwaal R, Wang J, Meijnen JP, et al. High-level production of beta-carotene in Saccharomyces cerevisiae by successive transformation with carotenogenic genes from Xanthophyllomyces dendrorhous. Appl Environ Microbiol. 2007;73(13):4342–4350. doi:10.1128/AEM.02759-06

36. Zhao Y, Zhang Y, Nielsen J, Liu Z. Production of β-carotene in Saccharomyces cerevisiae through altering yeast lipid metabolism. Biotechnol Bioeng. 2021;118(5):2043–2052. doi:10.1002/bit.27717

37. Jing Y, Guo F, Zhang S, et al. Recent Advances on Biological Synthesis of Lycopene by Using Industrial Yeast. Ind Eng Chem Res. 2021;60(9):3485–3494. doi:10.1021/acs.iecr.0c05228

38. Reyes LH, Kao KC. Growth-coupled carotenoids production using adaptive laboratory evolution. Methods Mol Biol. 2018;1671:319–330. doi:10.1007/978-1-4939-7295-1_20

39. Scita G. The stability of β-carotene under different laboratory conditions. J Nutr Biochem. 1992;3(3):124–128. doi:10.1016/0955-2863(92)90104-Q

40. Telfer A, De Las Rivas J, Barber J. β-Carotene within the isolated Photosystem II reaction centre: photooxidation and irreversible bleaching of this chromophore by oxidised P680. BBA - Bioenerg. 1991;1060(1):106–114. doi:10.1016/S0005-2728(05)80125-2

41. Bugaj LJ, Lim WA. High-throughput multicolor optogenetics in microwell plates. Nat Protoc. 2019;14(7):2205–2228. doi:10.1038/s41596-019-0178-y

42. Repina NA, McClave T, Johnson HJ, Bao X, Kane RS, Schaffer D V. Engineered Illumination Devices for Optogenetic Control of Cellular Signaling Dynamics. Cell Rep. 2020;31(10):107737. doi:10.1016/j.celrep.2020.107737

43. Heo J, Cho DH, Ramanan R, Oh HM, Kim HS. PhotoBiobox: A tablet sized, low-cost, high throughput photobioreactor for microalgal screening and culture optimization for growth, lipid content and CO2 sequestration. Biochem Eng J. 2015;103:193–197. doi:10.1016/j.bej.2015.07.013

44. Olson EJ, Hartsough LA, Landry BP, Shroff R, Tabor JJ. Characterizing bacterial gene circuit dynamics with optically programmed gene expression signals. Nat Methods. 2014;11(4):449–455. doi:10.1038/nmeth.2884

45. Wang H, Yang YT. Mini Photobioreactors for in Vivo Real-Time Characterization and Evolutionary Tuning of Bacterial Optogenetic Circuit. ACS Synth Biol. 2017;6(9):1793–1796. doi:10.1021/acssynbio.7b00091

46. Steel H, Habgood R, Kelly C, Papachristodoulou A. In situ characterisation and manipulation of biological systems with Chi.Bio. PLOS Biol. Published online 2020:1–12. doi:10.1371/journal.pbio.3000794

47. Nedbal J, Gao L, Suhling K. Bottom-illuminated orbital shaker for microalgae cultivation. HardwareX. 2020;8:e00143. doi:10.1016/j.ohx.2020.e00143

48. Pires JCM, Alvim-Ferraz MCM, Martins FG. Photobioreactor design for microalgae production through computational fluid dynamics: A review. Renew Sustain Energy Rev. 2017;79(May):248–254. doi:10.1016/j.rser.2017.05.064

49. Loomba V, Huber G, Von Lieres E. Single-cell computational analysis of light harvesting in a flat-panel photo-bioreactor. Biotechnol Biofuels. 2018;11(1):1–11. doi:10.1186/s13068-018-1147-3

50. Pfeiffer T, Morley A. An evolutionary perspective on the Crabtree effect. Front Mol Biosci. 2014;1(OCT):1–6. doi:10.3389/fmolb.2014.00017

51. Zhao EM, Lalwani MA, Chen J-M, Orillac P, Toettcher JE, Avalos JL. Optogenetic Amplification Circuits for Light-Induced Metabolic Control. ACS Synth Biol. 2021;10(5):1143–1154. doi:10.1021/acssynbio.0c00642

52. Zhao EM, Lalwani MA, Lovelett RJ, et al. Design and Characterization of Rapid Optogenetic Circuits for Dynamic Control in Yeast Metabolic Engineering. Published online 2020. doi:10.1021/acssynbio.0c00305

53. Benzinger D, Khammash M. Pulsatile inputs achieve tunable attenuation of gene expression variability and graded multi-gene regulation. Nat Commun. 2018;9(1). doi:10.1038/s41467-018-05882-2

54. Looser V, Bruhlmann B, Bumbak F, et al. Cultivation strategies to enhance productivity of Pichia pastoris: A review. Biotechnol Adv. 2014;33(6):1177–1193. doi:10.1016/j.biotechadv.2015.05.008

55. Aw R, Polizzi KM. Can too many copies spoil the broth? Microb Cell Fact. 2013;12(1):128. doi:10.1186/1475-2859-12-128

56. López J, Bustos D, Camilo C, Arenas N, Saa PA, Agosin E. Engineering Saccharomyces cerevisiae for the Overproduction of β-Ionone and Its Precursor β-Carotene. Front Bioeng Biotechnol. 2020;8(September):1–13. doi:10.3389/fbioe.2020.578793

57. Pouzet S, Banderas A, Bec M Le, Lautier T, Truan G, Hersen P. The promise of optogenetics for bioproduction: Dynamic control strategies and scale-up instruments. Bioengineering. 2020;7(4):1–17. doi:10.3390/bioengineering7040151

58. Gibson DG, Young L, Chuang R, et al. Enzymatic assembly of DNA molecules up to several hundred kilobases. 2009;6(5):12–16. doi:10.1038/NMETH.1318

59. Laughery MF, Hunter T, Brown A, et al. New vectors for simple and streamlined CRISPR-Cas9 genome editing in Saccharomyces cerevisiae. Yeast. 2015;32(12):711–720. doi:10.1002/yea.3098

60. Gietz RD, Schiestl RH. High-efficiency yeast transformation using the LiAc/SS carrier DNA/PEG method. Nat Protoc. 2007;2(1):31–34. doi:10.1038/nprot.2007.13

61. Ellis B, Haaland P, Hahne F, et al. flowCore: Basic structures for flow cytometry data. R Packag version 280. Published online 2022.

62. Llamosi A, Gonzalez-Vargas AM, Versari C, et al. What Population Reveals about Individual Cell Identity: Single-Cell Parameter Estimation of Models of Gene Expression in Yeast. PLoS Comput Biol. 2016;12(2):1–18. doi:10.1371/journal.pcbi.1004706

