## Supplementary Information for "Optogenetic control of beta-carotene bioproduction in yeast across multiple lab-scales"

In this document:

Supplementary Figure S1: Illumination quantifications across devices
Supplementary Figure S2: OptoTubes design
Supplementary Figure S3: OptoFlasks illumination stands designs
Supplementary Figure S4: Optogenetic activation timelines
Supplementary Figure S5: Lipid droplets microscopic observations
Supplementary Figure S6: Beta-carotene production timeline
Supplementary Figure S7: Yeast strains growth curves
Supplementary Figure S8: Beta-carotene pathway

Other files (available at https://github.com/Lab513/DIY_Optogenetics)

Supplementary File 1: OptoTubes STL Files
Supplementary File 2: eVOLVER design, hardware and software
Supplementary File 3: eVOLVER 3D-printed holder STL File

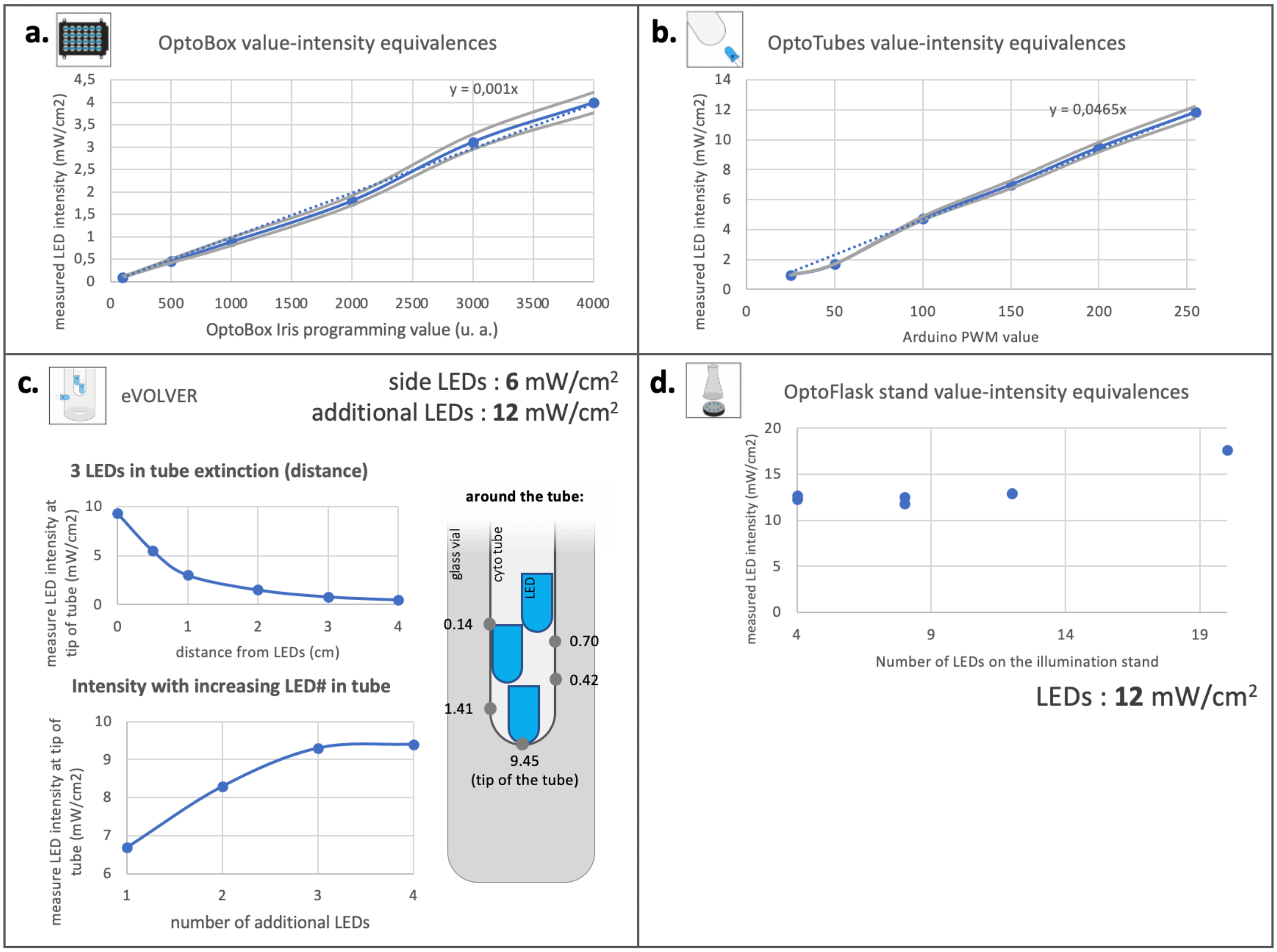


**Supplementary Figure S1 | Quantification of illumination across devices: set values (a.u.) *vs.* intensity (mW/cm$^2$). (a)** OptoBox equivalences were measured using a sensor placed directly above the LEDs, at the distance the imaging plate receives light. **(b)** The intensity of OptoTubes LEDs linearly increases with the PWM Arduino value up to a maximum of 12 mW/cm$^2$. Values were measured directly above (touching) the LED. **(c)** In the eVOLVER, intensity, and therefore illumination, decrease exponentially with the distance between the LED and the tip (base) of the tube containing the additional LEDs. Data for 3 additional LEDs are presented (top left). Bottom-left: increasing the number of LEDs in the tube increases the intensity but saturates at 3 LEDs. Right-side: scheme of the arrangement of the LEDs in the tube in the glass vial, and intensity values measured by sensor touching the tube at different locations around the tube. Note that to compute the total intensity in the medium, the 6 mW/cm$^2$ (the intensity set by a resistor) of the side-LED should be added to these measures. **(c)** Measured intensity in the OptoFlask illumination stand: all LEDs display 12 mW/cm$^2$. The higher value observed for 20 LEDs is due to light leaking from nearby LEDs. Measures were taken on top of the petri-dish covering the LEDs.

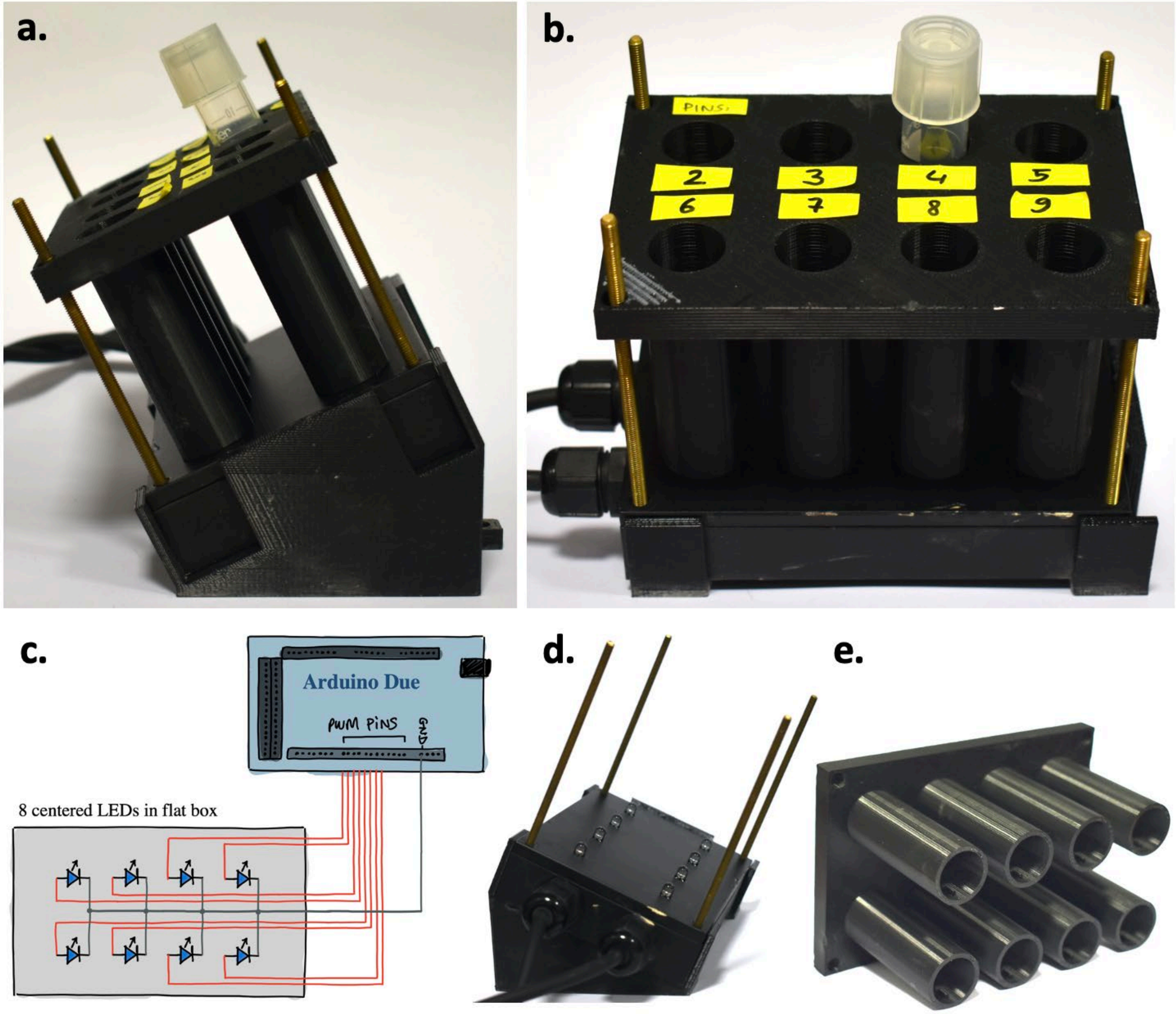


**Supplementary Figure S2 | Design of the OptoTubes.** **(a)** Side-view and **(b)** front view of the device. **(c)** Electronic connection inside the flat box containing the LEDs, positioned on its 3D-printed stand in **(d)**. **(e)** 3D-printed tube holder (which holds and positions the culture tubes above the LEDs). Long screws are used to secure the entire structure. The STL files for the 3D-printed parts are available in Supplementary File 1.

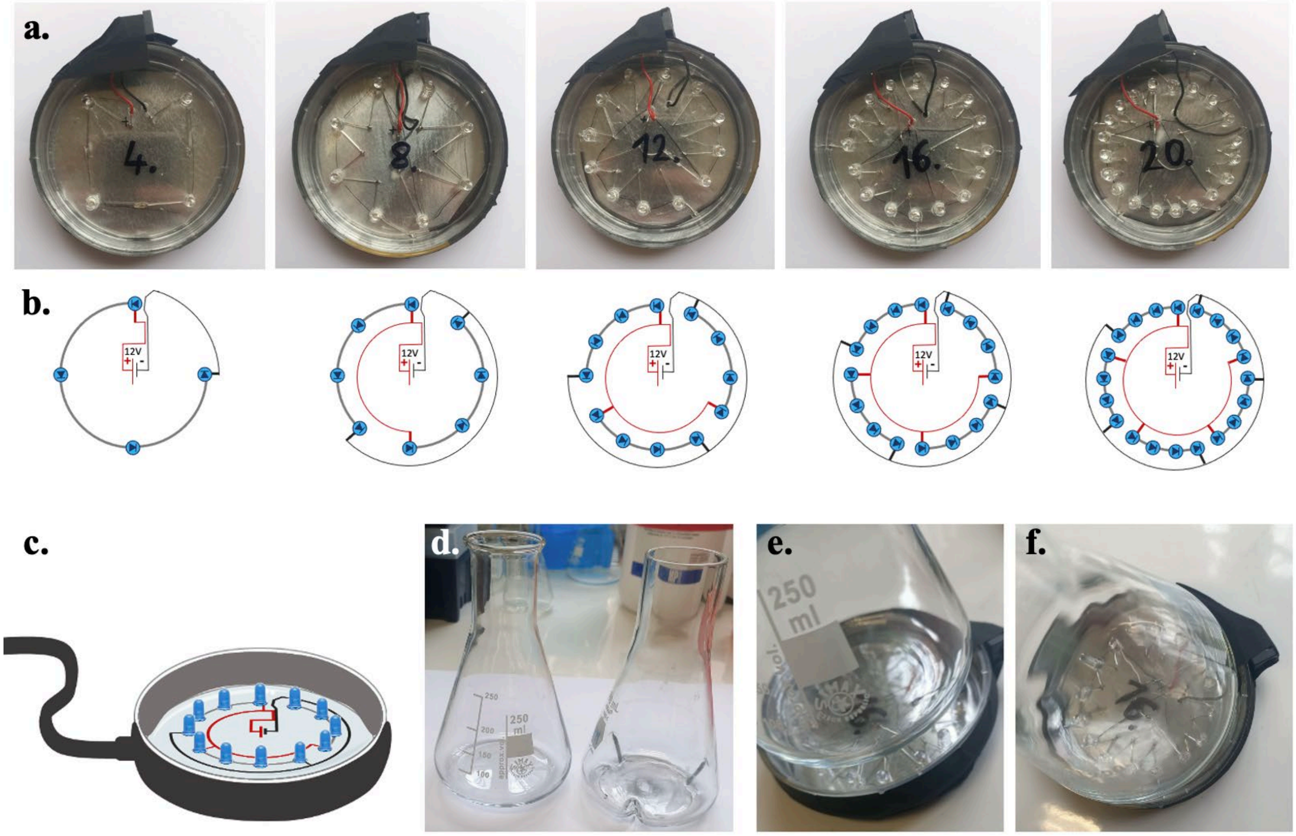


**Supplementary Figure S3 | Designs of OptoFlasks illumination stands**. **(a)** Pictures of the five types of illumination stands with 4, 8, 12, 16 and 20 LEDs. The hump on top corresponds to the DC power supply jack connector (12V). **(b)** Corresponding electrical designs that fit inside Petri dishes and ensure same light intensities between LEDs and between illumination stands. **(c)** Side-view scheme of the illumination stand in which 250 mL flasks, either flat or indented/baffled **(d)** are placed. **(e., f.)** To maximize illumination, the circular pattern of LEDs lines up with the edges of the flask, where the medium will localize in a shaking incubator.

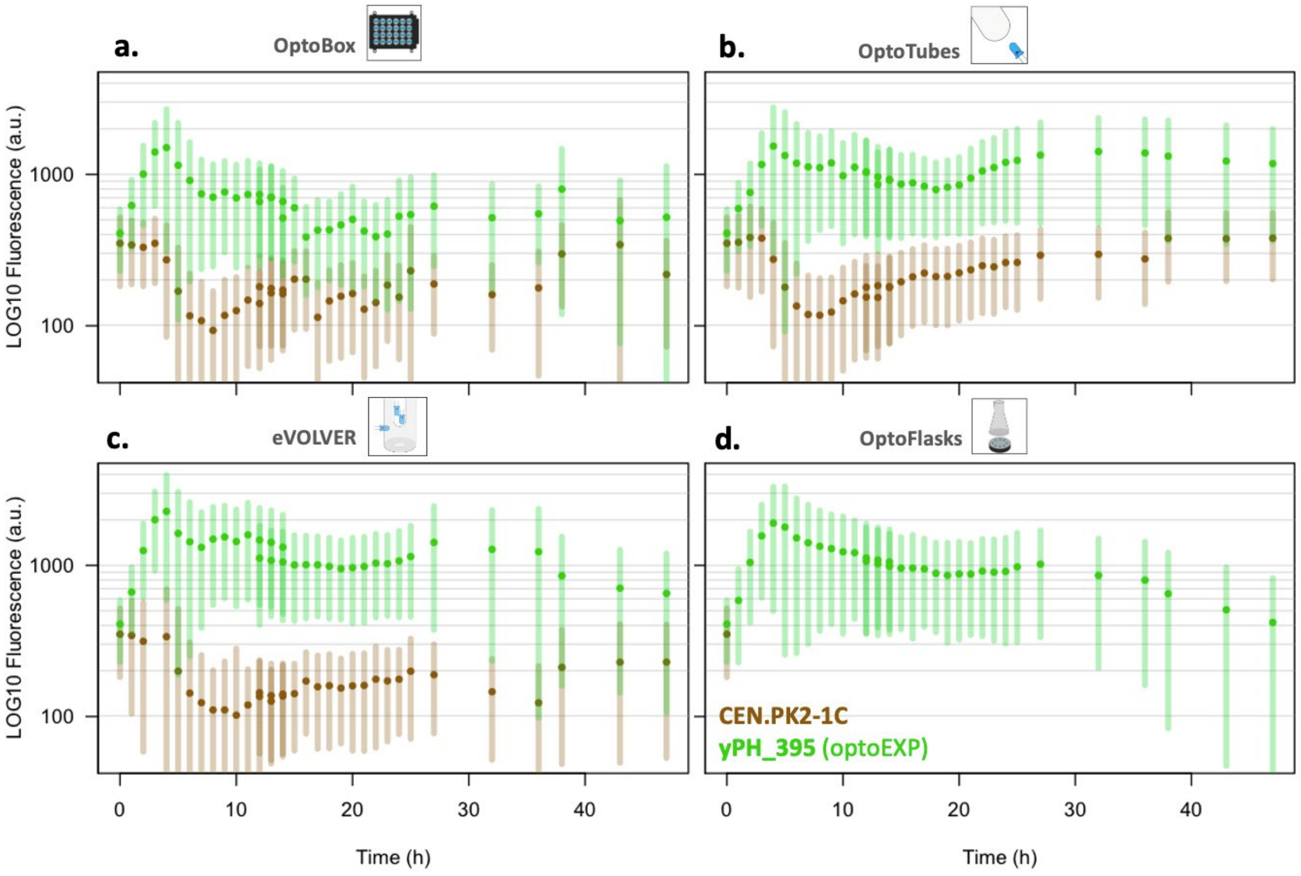


**Supplementary Figure S4 | Timelines of optogenetic activation in the different devices**. Brown points are the CEN.PK2-1C WT strain. Green points are the OPTO-EXP strain. Error bars represent the standard deviation fluorescence value of the 10 000 cells quantified. The exposure to light of a culture with an $OD_{600}$ 0.05 starts at T0; GFP level was measured every hour up to 25 hours, then at 27, 32, 36, 38, 43 and 47 hours. **(a)** Activation in the OptoBox, 1 mL cultures, illumination is 8000 (max). **(b)** Activation in the OptoTubes: 3 mL cultures, illumination 255 (max). **(c)** Activation in eVOLVER: 15 mL cultures, stirring 255, s+2 illumination. **(d)** Activation in OptoFlasks: 50 mL cultures in 250 mL indented flasks, 12 LED illumination stand.

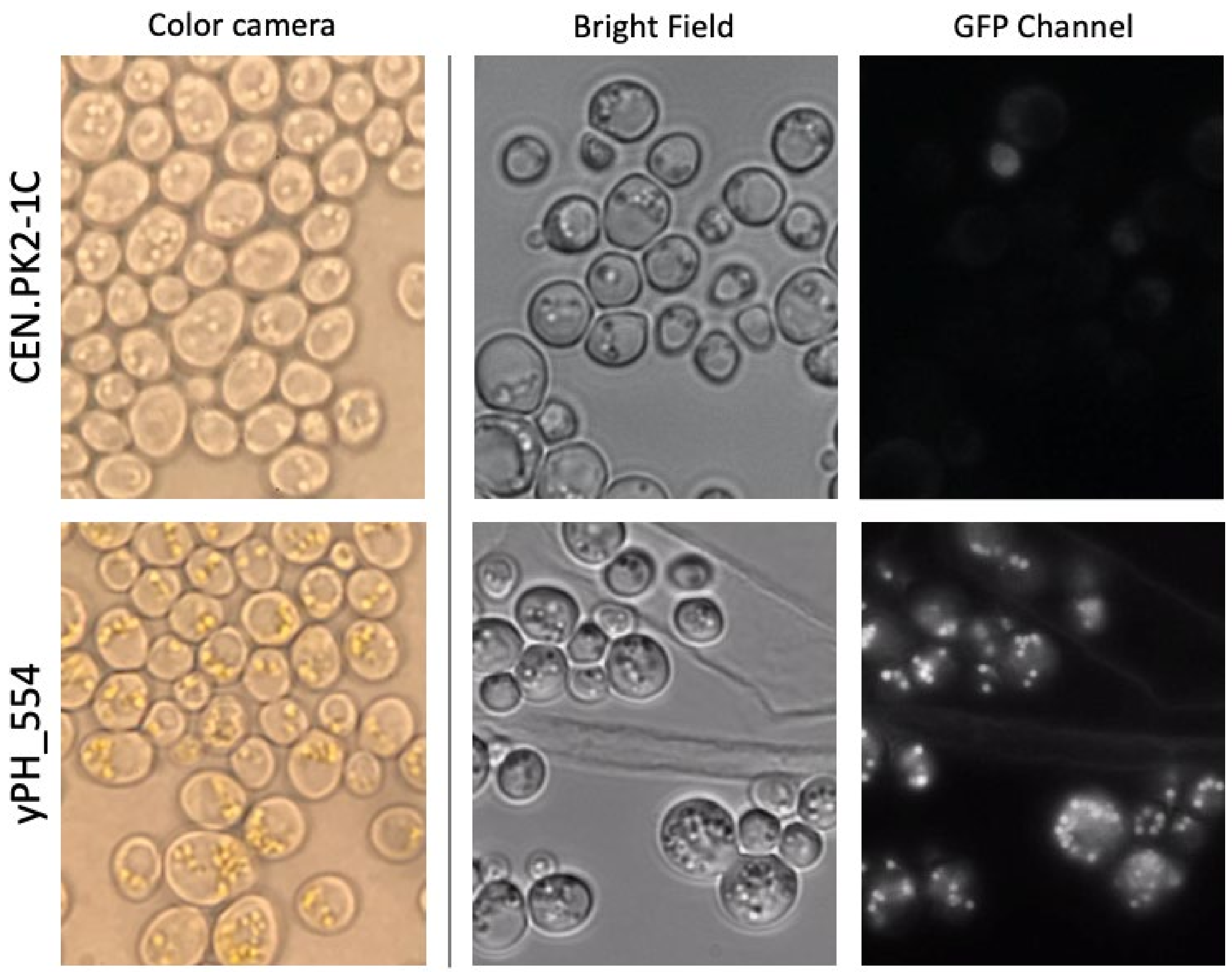


**Supplementary Figure S5 | Microscopic observations of lipid droplets (100X).** Beta-carotene localizes in lipid droplets. **(Left)** Color camera view (does not correspond to the other microscopic pictures). Note the golden droplets are only present in the cells of the constitutive producer strain yPH_554. It is necessary to adjust with the microscope diaphragms to obtain a lighting setting that display the yellow color, besides using a color camera (these images were taken with a phone through an ocular lens). **(Middle)** Bright field (exposure 1 sec) image and corresponding fluorescent (exposure 1 sec) image **(Right)**.

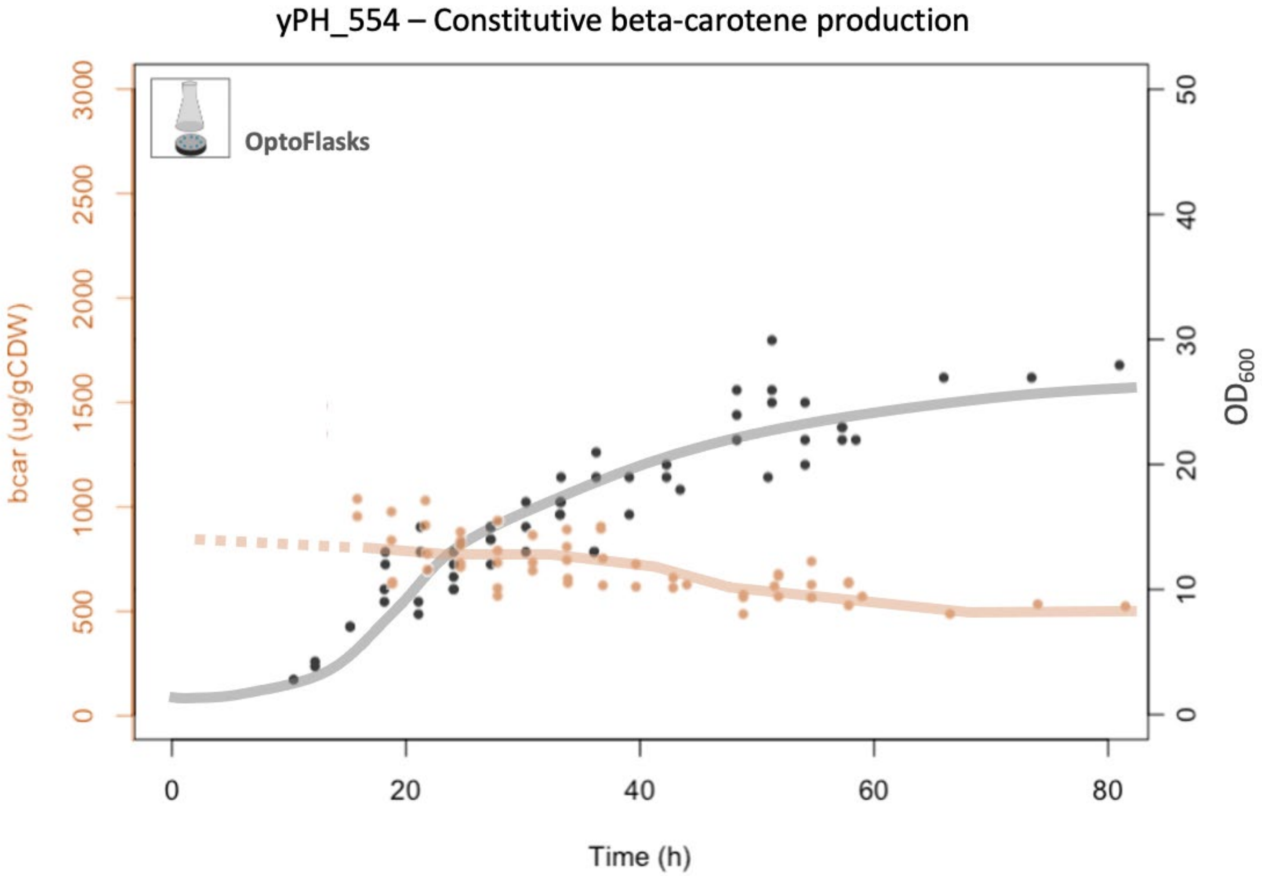


**Supplementary Figure S6 | Beta-carotene production during cultivation of strain yPH_554**. Each point corresponds to one measure, beta-carotene content was estimated as described in the materials and methods. $OD_{600}$ was measured by hand using a spectrophotometer. The small number of cells at the beginning of the culture prevented early quantification of beta-carotene (dashed line). The hand-drawn solid lines illustrate proposed trends – the grey line is estimated from Figure S7. Growth and production in the dark, 50 mL culture in a 250 mL indented flask; 24 h was chosen as appropriate timepoint to estimate beta-carotene production under different conditions.

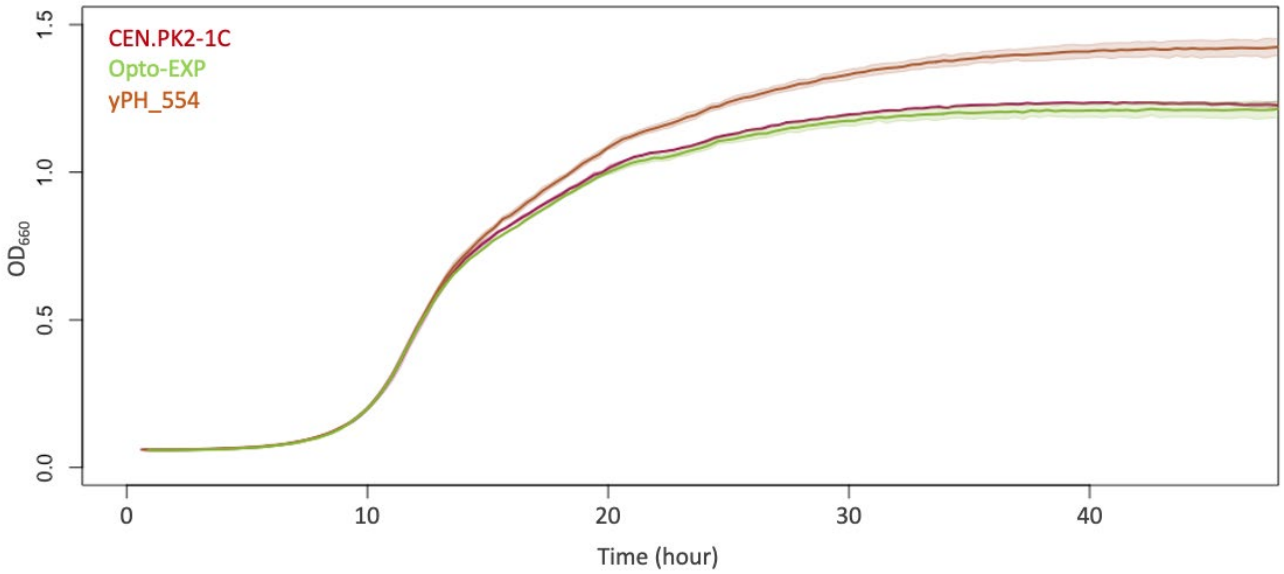


**Supplementary Figure S7 | Yeast strain growth curves.** Cultures were performed in filter sterilized YPD and growth was quantified by reading optical density values using a plate reader. No burden was detected for the beta-carotene constitutive producer strain yPH_554 compared to the non-producer strains, i.e., the control CEN.PK2-1C and the optogenetic strain OPTO-EXP: the exponential growth phase was not impacted, and the other phases of growth appear to lead to increased total biomass, perhaps due to lower overflow metabolism in the fermentation phase. *N*=2.

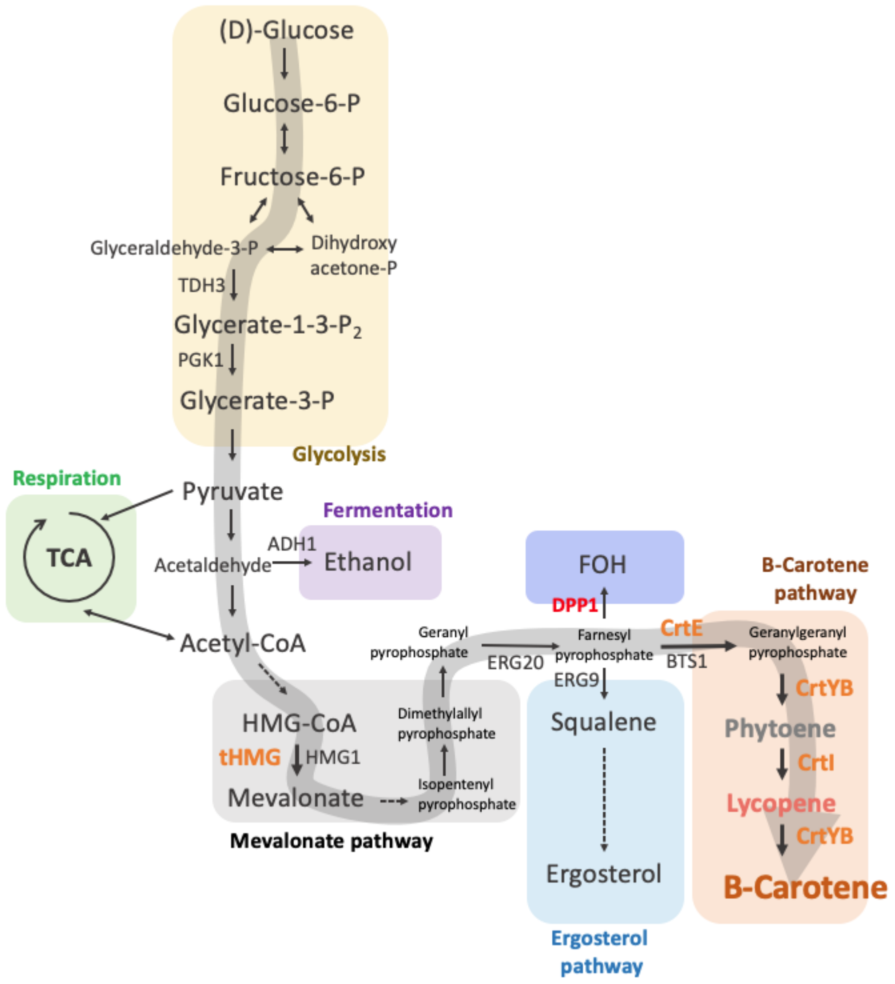


**Supplementary Figure S8 | Yeast metabolic pathways.** With detailed chemical names.
